# Runs of Homozygosity in sub-Saharan African populations provide insights into a complex demographic and health history

**DOI:** 10.1101/470583

**Authors:** Francisco C. Ceballos, Scott Hazelhurst, Michele Ramsay

## Abstract

The study of runs of homozygosity (ROH), contiguous regions in the genome where an individual is homozygous across all sites, can shed light on the demographic history and cultural practices. We present a fine-scale ROH analysis of 1679 individuals from 28 sub-Saharan African (SSA) populations along with 1384 individuals from 17 world-wide populations. Using high-density SNP coverage, we could accurately obtain ROH as low as 300Kb using PLINK software. The analyses showed a heterogeneous distribution of autozygosity across SSA, revealing a complex demographic history. They highlight differences between African groups and can differentiate between the impact of consanguineous practices (e.g. among the Somali) and endogamy (e.g. among several Khoe-San groups^1^). The genomic distribution of ROH was analysed through the identification of ROH islands and regions of heterozygosity (RHZ). These homozygosity cold and hotspots harbour multiple protein coding genes. Studying ROH therefore not only sheds light on population history, but can also be used to study genetic variation related to the health of extant populations.

## INTRODUCTION

African human genetic diversity provides the ideal backdrop to reconstruct modern human origins, the genetic basis of adaptation to different environments and the development of more effective vaccines^1^. Studies on African population genetics and genomics have multiplied over the past decade, boosted by many efforts to genotype and sequence more populations from the continent^2-4^, though one of the “grand challenges” of the post-genome era, “To characterize genetic variation among individuals and populations”^5^, is yet to be fully achieved. Testament to the value of this approach is the recent study of the deep whole genome sequencing of 24 South African individuals where roughly 0.8M new variants were identified^6^. Due to the significant advances in genotyping and sampling of African populations, a study on runs of homozygosity provides an interesting opportunity for a deep dive into the demographic history of Africans.

Runs of homozygosity (ROH) are contiguous regions of the genome where an individual is homozygous (autozygous) across all sites^7^. ROH arise when two copies of an ancestral haplotype are brought together in an individual. The size of the ROH is inversely correlated with its age: longer ROH will be inherited from recent common ancestors while shorter ROH from distant ancestors because they have been broken down by recombination over many generations. Very short ROH, characterized by strong linkage disequilibrium (LD) among markers, are not always considered autozygous but nevertheless are due to the mating of distantly related individuals. A different source of apparent homozygosity, hemizygous deletions, can masquerade as ROH, but such copy number variation has a minor effect in ROH studies^7-9^.

Since their discovery in the mid-1990s^10^ ROH were found to be ubiquitous. We are all inbred to some degree and ROH capture this aspect of our demographic histories, with runs of homozygosity being the genomic footprint of the phenomenon known as pedigree collapse^11^. ROH are present in all populations, even in admixed or outbred populations and arise by two different processes: a limited effective population size (Ne) and by consanguineous unions. Independently of how they were generated, ROH can be used to obtain the genomic inbreeding coefficient or F_ROH_^7; 8^. Traditionally, the inbreeding coefficient (the probability that an individual receives two alleles that are identical-by-descent at a given locus which is also the expected proportion of the genome being autozygous) is obtained using pedigrees and its accuracy depends on the depth and reliability of the pedigree^12; 13^. F_ROH_ measures the actual proportion of the autosomal genome that is autozygous over and above a specific minimum length ROH threshold. When this cut-off is set at 1.5Mb, F_ROH_ correlates most strongly (r=0.86) with the F obtained from an accurate six-generation pedigree (F_PED_)^8^. Using 20-generation depth genealogies with more than 5000 individuals of European Royal dynasties, with many complex inbreeding loops, it has been found that above the 10^th^ generation the change in the coefficient of inbreeding (F) is less than 1%^14^. Also, it has been found that individuals with no inbreeding loops in at least 5 generations (and probably 10) carried ROH up to 4Mb in length but not longer^8^. F_ROH_, using a genomic approach, captures the total inbreeding coefficient of the individual independently of pedigree accuracy, or depth within the resolution of the data available and the size of ROH that can be called^7; 15^.

The ROH approach provides a window to explore individual and demographic history, to understand the genetic architecture of traits and diseases and to study concepts in genome biology^7^. Different population histories give rise to divergent distributions of long and short ROH. The number and length of ROH reflect individual and population history and have been used to detect consanguineous practices, endogamy and isolation^7; 9^. ROH were found to be associated with different diseases and traits and its analysis is capable of detecting directional dominance and inbreeding depression when phenotype data are available^16; 17^.

The non-random patterns of the genomic distribution of ROH provides an interesting approach to studying genome biology^7; 18-20^. As expected, ROH are common in regions of high LD, low recombination and low genetic diversity^19; 20^. There is an uneven distribution along the genome, with a number of comparatively short regions with a high population-specific prevalence of ROH – known as ROH islands – on each chromosome, as well as coldspots with a paucity of ROH^20; 21^. These ROH islands are prevalent in all populations and dominate the ROH in outbred groups; however they are overshadowed by much larger ROH arising from recent pedigree loops that are randomly distributed across the genome^7^. In some cases, ROH islands are due to homozygosity of one common haplotype, but in other cases, multiple haplotypes contribute to a single ROH island^20^. The origin of these islands is still a subject of debate. In some cases, the haplotypes segregating at high frequencies in the population may be due to positive selection; for example, a ROH island around the lactase persistence (*LCT*) gene on chromosome 2q21 was found in Europeans^21^. In addition, numerous genes that are targets of recent positive selection have been found in multiple ROH islands in populations around the globe^20^. Another potential biological explanation is that ROH islands include small inversions that suppress recombination^21^.

Sub-Saharan Africa (SSA) is a sub-continent with a complex demographic history where a deep ROH analysis provides interesting insights. Previous studies on ROH were hampered by small sample sizes and inadequate African population representation, genotype panels with low SNP coverage, non-optimized ROH calling conditions and in some cases poor ROH classification and analysis. Gibson et al.^18^, in one of the first articles that included African samples, published in 2006, used the Hap Map I dataset with 60 Yoruba individuals to conclude that Western Africans had the smallest number of long ROH tracks per individual, but showed that ROH are common even in outbred populations. Four years later, Kirin et al.^9^ used the Human Genome Diversity Project to analyse five SSA populations: three agricultural heritage and two hunter-gatherer groups with 82 individuals in total. With a panel of 415K SNPs the study concluded that populations in SSA have the fewest ROH, for any ROH size, in comparison to other world populations, and that there is an increase in ROH with distance from Africa. The article also suggested that the hunter-gatherers (17 Biaka and Mbuti pygmies and 15 !Xun San) have a larger ROH burden between 0.5 and 16Mb compared to farmer communities. Henn et al.^22^ used 90 hunter gatherer individuals from three populations (Hadza, Sandawe and ≠Komani) to calculate the cumulative ROH (cROH) as the sum of ROH >500kb. They concluded that the Hadza population differ strongly from the other groups and its elevated mean and variance of cROH is indicative of a severe population bottleneck. Further evidence of the heterogeneity among the hunter-gathered populations from SSA was reported by Schlebusch et al.^23^. Using a sliding window of 5Mb and a coverage of 297K SNPs, a minimum length of 500kb and 50kb/SNP in PLINK they obtained the cROH for 147 individuals from 21 populations (9 farmers and 12 hunter-gatherer populations). Considering the heterogeneity among hunter-gatherers the study concluded that northern San groups like /Gui and //Gana, Nama and the two Pygmy populations have generally an average cROH higher than farming populations for every ROH size class. However, southern San groups (Karretjie and ≠Khomani) have a lower burden than farmers. In one of the first studies to provide a meaningful world context of the distribution of ROH, Pemberton et al.^20^ analysed 64 worldwide populations (1839 individuals in total) including 10 from SSA (2 hunter-gatherer and 8 farmer-pastoralist populations (386 individuals in total)). After identifying ROH by a LOD score methodology, and using a mixture of three Gaussian distributions, ROH were classified by length into 3 groups: Class A (short ROH of about tens of kb with an LD origin), Class B (intermediate ROH of hundreds of kb to 2 Mb, resulting from background relatedness owing to genetic drift) and Class C (long ROH over 1 – 2 Mb arising from recent parental relatedness). The study concluded that Class A and B ROH increase with distance from Africa, a trend similar to the negative correlation observed for expected heterozygosity^24^. Class C ROH did not show this geographical stepwise increase; however, African populations tended to have few ROH in this class.

Representation of SSA populations has increased with projects such as the AGVP^2^, 1000 Genomes Project^25^, the HGDP^26^, the Simons Genome Diversity Project^27^, and others^22; 23; 26; 28^, making it possible to study 3000 individuals in over 60 SSA populations. Recent studies have, however, shown that the distribution of ROH in SSA may not be as homogeneous as previously thought. Hollfelder et al.^29^ genotyped 244 new individuals from 18 Sudanese populations and, notwithstanding some technical issues, concluded that Coptic, Cushitic, Nubian and Arabic populations from North Sudan have a higher burden of ROH in comparison to Southern Sudan populations. ROH distribution heterogeneity in SSA was also shown by Choudhury et al.^6^ by analysing roughly 1600 individuals from 28 SSA populations, in a preliminary superficial exploration. Finally, Ceballos et al.^7^ gathered more than 4200 individuals from 176 worldwide populations to analyse ROH distribution. Although this study included 924 SSA individuals from 30 population, the low SNP coverage (147K SNPs) prevented fine-scale analysis, but concluded that some hunter gatherer populations like the Hadza have a ROH burden similar to the most isolated populations from Oceania and South America.

The objective of this study was to perform fine-scale analysis of the ROH distribution in SSA, in a world context, in order to learn more about the demographic history of the continent and its populations. Public data from the Africa Genome Variation Project (AGVP), the 1000 Genome Project (KGP) and Schlebusch et al. were analysed and included 1679 individuals from 28 SSA populations and 1384 individuals from 17 worldwide populations. By analysing the sum and number of ROH and deconstructing probable patterns of inbreeding, we present interpretations for the demographic histories of different SSA populations.

## Materials and Methods

### Description of the Data

The study included a total of 3063 individuals from 45 populations from the 1000 Genomes Project – Phase 3 (KGP)^25; 30^, the African Genome Variation Project (AGVP)^2^ and Schlebusch et al. (2012)^23^. All individuals were genotyped using the Infinium Omni 2.5 array from Illumina, and all datasets were subjected to extensive QC procedures.

The KGP – Phase 3, includes a total of 1558 individuals from 19 populations^25^. From Europe: FIN (Finish in Finland, n=97), GBR (British in England and Scotland, n=91), IBS (Iberian populations in Spain, n=99), TSI (Tuscany in Italy, n=92) and CEU (Utah residents with European ancestry=95). From America: ASW (Americans of African ancestry in Houston, n=49), ACB (African Caribbean in Barbados, n=72), PUR (Puerto Rican in Puerto Rico with admixed ancestry, n=72), PEL (Peruvian in Lima, Peru with Amerindian ancestry, n=50), CLM (Colombian in Medellin, Colombia with admix ancestry, n=65) and MXL (Mexican with admixed ancestry in Los Angles, USA, n=47). From South Asia: GIH (Gujarati Indian from Houston, Texas n=95). From East Asia: CDX (Chinese Han in Xishuangbanna, China, n=83), CHB (Chinese Han in Beijing, China, n=98), CHS (Southern Han Chinese, n=86), JPT (Japanese in Tokyo, Japan, n=96) and KHV (Kinh in Ho Chi Minh city, Vietnam n=96). From Africa Guinean Gulf: YRI (Yoruba in Ibadan, Nigeria, n=100), and from East Africa: LWK (Luhya in Webuye, Kenya, n=74).

The AGVP includes 1318 individuals from 17 populations from SSA^2^. Niger-Congo speakers from Western Africa: Wolof (Senegambian sub-group speakers from The Gambia, n=78), Fula (Senegambian from The Gambia, n=74), Mandinka (Mande sub-group speakers from The Gambia, n=88) and Jola (Bak sub-group speakers from The Gambia, n=79). Niger-Congo speakers from the Guinean Gulf: Ga-Adangbe (Kwa sub-group speakers from Ghana, n=100) and Igbo (Igboid sub-group speakers from Nigeria, n=99). Afro-Asiatic speakers from the Horn of Africa: Amhara (Semitic sub-group speakers from Ethiopia, n=42), Oromo (Cushitic sub-group speakers from Ethiopia,n=26) and Somali (Cushitic from Ethiopia and Somalia, n=39). Niger-Congo speakers from Eastern Africa: Baganda (Bantoid sub-group speakers from Uganda, n=100), Banyarwanda (Bantoid from Uganda, n=100), Barundi (Bantoid from Uganda, n=97) and Kikuyu (Bantoid from Kenya, n=99). Nilo-Saharan speakers from Eastern Africa: Kalenjin (Eastern Sudanic sub-group speakers from Kenya, n=100). Niger-Congo speakers from Southern Africa: Sotho (Bantoid from South Africa, n=86) and Zulu (Bantoid from South Africa, n=100).

In addition, 147 individuals from 7 different groups with Khoe and San ancestry, 40 South African Colored individuals (20 from Colesberg and 20 from Wellington, both in South Africa) and 12 Herero Bantoid speakers from Namibia from the Schlebusch study were added ^23^. The term Khoe-San designates two groups of people: the pastoralist Khoe and the hunter-gatherer San^23; 31^. The following were included in this study: Ju/’hoansi (San Ju speakers from Namibia, n=18), !Xun (San Ju speakers Angola, n=19), Gui//Gana (San Khoe-Kwadi speakers from Botswana, n=15), ≠Khomani (San Tuu speakers from South Africa, n=39), Nama (Khoe Khoe-Kwadi speakers from Namibia), Khwe (San Khoe-Kwadi speakers from the Caprivi strip: Namibia, Angola and Botswana) and Karretjie people (San Tuu speakers from South Africa, n=20).

SSA samples were grouped according to geographic region and principal components analysis into 8 groups (Figure 1): Western Africa (n=319), Gulf of Guinea (n=299), Eastern Africa Niger-Congo populations (n=470), Eastern Nilo-Saharan population (n=100), Horn of Africa (n=107), Southern Africa (n=198), Khoe and San groups (n=147) and Colored South Africans (n=40). KGP populations from the rest of the world were grouped as follows: Mixed African-American populations (n=121), Europeans (n=474), Southern Asians (n=95), Eastern Asians (n=459), South Americans (n=50) and Mixed Hispanic-Americans (n=184). Since the three datasets used in this study were genotyped using the same SNP genotyping array they could easily be merged ^15; 16^. Only autosomal SNPs were included in this analysis. For each population, array data were filtered to remove SNPs with minor allele frequencies < 0.05 and those that divert from H-W proportions with *p* < 0.001. This filtering serves to limit the effects of ascertainment bias caused by the small number of individuals in the SNP discovery panel. After QC, there were 1.3M SNPs on average in Western Africa populations, 1.4M in Gulf of Guinea, 1.4M in Eastern Africa Niger-Congo populations, 1.4M in Eastern Nilo-Saharan population, 1.3M in Horn of Africa populations, 1.3M in Bantu-speaking Southern Africa populations, 1.4M in Khoe and San populations from Southern Africa, 1.4M in Colored populations from Southern Africa, 1.4M in Africa-American admixed populations, 1.2M in European populations, 1.2M in southern Asia populations, 1.1M in Eastern Asian populations, 1.1M in South America populations and 1.2M in Hispanic-American admixed populations.

**Figure 1.**
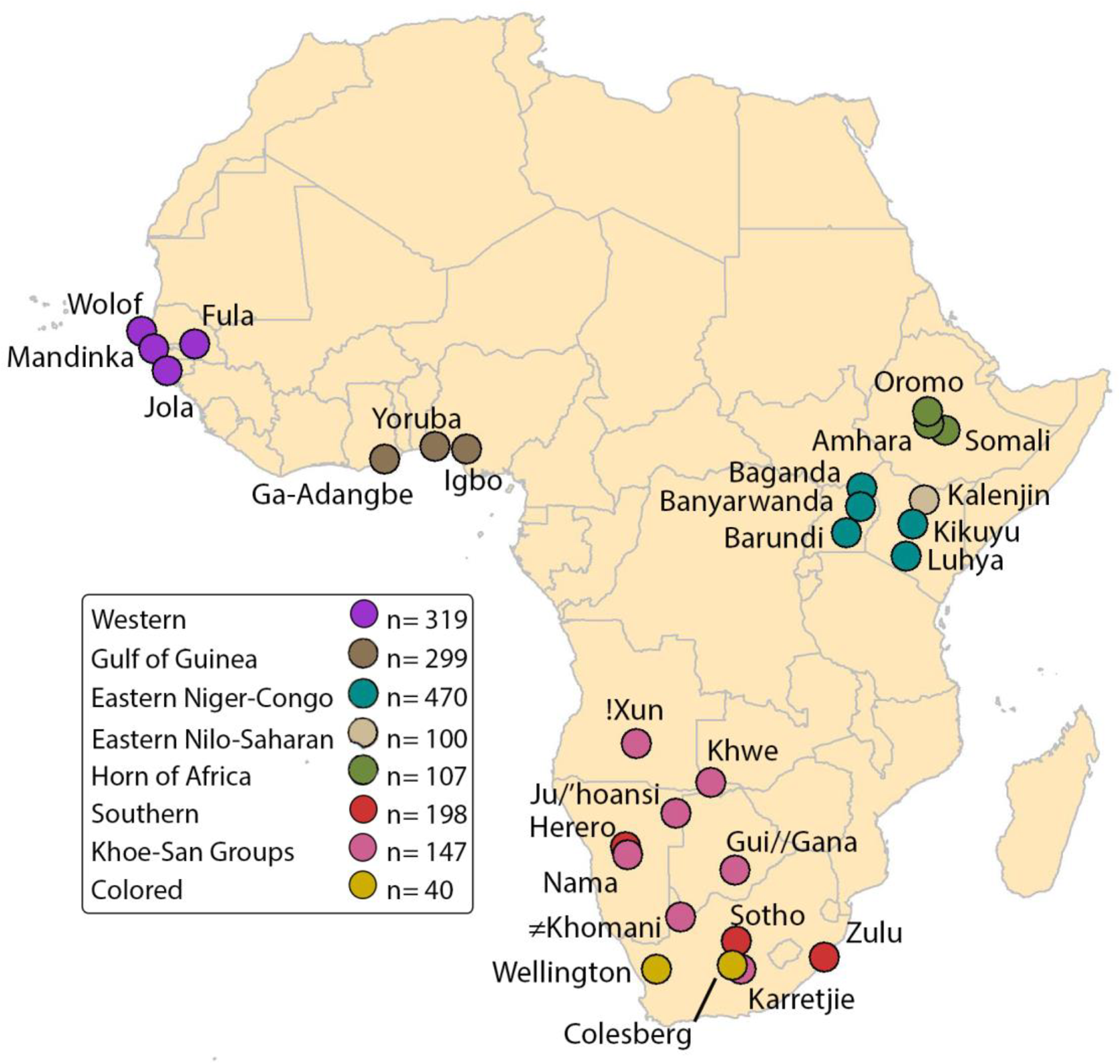
Sub-Saharan African populations included in the study: 28 African populations in total including 16 from the African Genome Variation Project (AGVP), 2 from the 1000 Genomes Project (KGP) and 10 from Schlesbusch et al. 2012. Populations were organized in 8 groups according to their geographic, linguistic and/or admixture origins. Western Africa (shown in deep purple), Gulf of Guinea (shown in brown), Eastern Africa Niger-Congo populations (shown in light blue), Eastern Africa Nilo-Saharan population (shown in wheat), Horn of Africa (shown in dark green), Southern Africa (shown in red), Khoe and San populations (shown in pink) and Colored admixed populations (shown in yellow). The number of individuals from each group is shown in Table 1.g

### Merging with the Human Genome Diversity Project Data

To enrich the data further we merged the above datasets (KGP, AGVP and Schlebusch) with the Human Genome Diversity Project dataset (HGDP)^26^ since this dataset includes isolates and urban populations from across four continents. The HGDP includes 1043 individuals from 51 populations from different parts of the world: 6 populations from Europe, 4 from the Middle East, 10 from Central and South Asia, 17 from East Asia, 7 from Africa, 2 from Oceania and 5 from Africa. 650K SNPs were genotyped in these populations using the Illumina BeadStation technology. After merging all datasets and filtering for MAF and H-W proportions we have a dataset of 4106 individuals with genotypes for 382,840 SNPs. In order to differentiate it from the main dataset described above, this merged dataset is called “*worldata0.3*”.

### Identification of runs of homozygosity

The observational approach implemented by PLINK v1.9^32^ was used to call ROH. The simplicity of the approach used by PLINK allows efficient execution on data from large consortia and even different array platforms or sequencing technologies^7; 16^. Tests on simulated and real data showed that the approach used by PLINK outperformed its competitors in reliably detecting ROH^33^.

The following PLINK conditions were applied to search for ROH:

`--homozyg-snp` 30. Minimum number of SNPs that a ROH is required to have

`--homozyg-kb 300.` Length in Kb of the sliding window

`--homozyg-density 30.` Required minimum density to consider a ROH (1 SNP in 30 Kb)

`--homozyg-window-snp 30.` Number of SNPs that the sliding window must have

`--homozyg-gap 1000.` Length in Kb between two SNPs in order to be considered in two different segments.

`--homozyg-window-het 1.` Number of heterozygous SNPs allowed in a window

`--homozyg-window-missing 5.` Number of missing calls allowed in a window

`--homozyg-window-threshold 0.05.` Proportion of overlapping window that must be called homozygous to define a given SNP as in a “homozygous” segment.

The objective of this study is to use autozygosity to learn more about demographic history in SSA populations. To achieve this goal short and long ROH need to be explored, since they provide different types of information^7; 15^. The high SNP coverage of 1.2M SNPs on average for all the populations included in the study, makes it possible to find a single SNP, on average, in a track of 2.4 Kb. The Supplemental Methods and Figures S1, S2, S3, S4 and S5 demonstrate that this coverage allows accurate detection of ROH longer than 300 Kb by considering 30 as a minimum number of SNPs per ROH and/or the required minimum SNP density to call ROH. To obtain a window with 30 SNPs, on average (assuming a homogeneous distribution of SNP along the genome), a tract of just 72 Kb is needed. A threshold of 300 Kb was set for the minimum length in order to capture small ROH originating far in the past and also to ensure that these are true ROH that originated by genetic drift or consanguinity. An alternative source of homozygosity originating from linkage disequilibrium (LD) typically produces tracts measuring up to about 100 Kb, based on empirical studies^34-36^. By using a minimum-length cutoff of 300 Kb, most short ROH resulting from LD will be eliminated.

### Analyses

Different variables were obtained and analyses performed in order to fully exploit the usefulness of the ROH in the understanding of demographic history and possible cultural practices of populations. First, we obtained the total sum of ROH for six ROH length classes: 0.3 – 0.5, 0.5 – 1, 1 – 2, 2 – 4, 4 – 8 and >8 Mb. This exploratory data analysis allows us to delve into aspects of population history, since, due to recombination, the size of a ROH is inversely proportional to its age. Thus, plotting the total sum of ROH for these size classes will inform, for example, the relative change of the effective population size across generations.

We also conducted a preliminary examination at a global level using *worldata.03*. The interest in this exploratory data analysis is to provide a rough relative comparison among populations not an absolute quantification, as the lower SNP density affects the accuracy of analysis (it is apparent in Figure S6 that very short and large ROH are underestimated in *worldata.03* due to the lower SNP coverage, and the degree of bias depends on the population and its genetic characteristics). However, in further analysis, where absolute quantification and comparison is mandatory in order to obtain meaningful conclusions, the underestimation of short and very long ROH prevents the use of *worldata.03*.

For comparison purposes four variables were defined: (1) *Mean number of ROH* as the population average number of ROH longer than 1.5 Mb; (2) *Mean ROH size* as the population average size of ROH longer than 1.5Mb; (3) *Total sum of ROH>1.5 Mb* as the population average total sum of ROH longer than 1.5 Mb; and (4) *Total sum of ROH<1.5* as the population average total sum of ROH shorter than 1.5 Mb. Exploratory data analysis and data representation were illustrated using violin plots. These plots combine a box plot with a kernel density plot, where the interval width is obtained by the rule of thumb. The violin plot shows a colored density trace with the interquartile range as a black line and median as a white dot. This representation is especially useful when dealing with asymmetric distributions where median is more informative than the mean. Statistical comparisons between total sum of ROH longer and shorter than 1.5 Mb between populations and geographic regions were performed using the Whitney-Wilcoxon non-parametrical test (MWW). All the analyses were performed using R (v.3.4.1)^37^.

### Measuring different sources of inbreeding

Population geneticists use the word inbreeding to mean different things, as pointed out by Jacquard and Templeton in their respective classic articles^38; 39^. Inbreeding can be produced by a deviation from panmixia, in what G. Malecot called systematic inbreeding, or by genetic drift and low effective population size, also called panmictic inbreeding^40^. Systematic inbreeding has a direct effect on the H-W proportions of a population and can be measured using the Wright’s fixation index or F_IS_^41^. In this study this component of the total inbreeding coefficient is measured using the --het function in PLINK. In this context F_IS_ is the average SNP homozygosity within an individual relative to the expected homozygosity of alleles randomly drawn from the population. PLINK use the following expression:

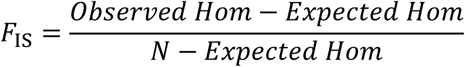

where *Observed Hom* is the observed number of homozygous SNPs, *Expected Hom* is the expected number of homozygous SNPs considering H-W proportions and *N* is the total number of non-missing genotyped SNPs. F_IS_ thus measures inbreeding in the current generation with F_IS_ = 0 indicating random mating, F_IS_ > 0 indicating consanguinity and F_IS_ < 0 indicating inbreeding avoidance.

The two different sources of inbreeding, namely, genetic drift (denoted by F_ST_) and non-random mating (F_IS_) are both components of the total inbreeding coefficient (F_IT_), defined as the probability than an individual receives two alleles that are identical-by-descent. Sewall Wright developed an approach to consider these three different F coefficients in his F statistics (1-F_IT_)=(1-F_IS_)(1-F_ST_)^41; 42^. First defined as correlations, Nei showed how these coefficients can be expressed in terms of allele frequencies and observed and expected genotype frequencies^43^. In this framework, F_ST_ can be considered a measure of the genetic differentiation of a subpopulation in comparison with an ideal population with a large N_e_. F_IT_ is the total inbreeding coefficient, traditionally obtained using deep genealogies, and can be calculated using the F_ROH_ (ROH > 1.5Mb):

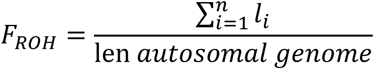

Where the numerator is the sum of n ROH of length l_i_ (>1.5Mb) and the denominator is the total autosomal length.

### Genomic distribution of ROH

The study of the genomic distribution of ROH can be used for different purposes. By identifying the regions where ROH are very prevalent, or completely absent in the population it is possible to identify candidate regions (including protein coding genes) under selection. Furthermore, the identification of common and unique ROHi in the different regional groups considered in this study can also shed light on population demographic history. In order to study the spatial distribution of ROH across the genome two different variables were defined: islands of runs of homozygosity (ROHi) and regions of heterozygosity (RHZ) (see definitions below). In order to identify protein coding genes in these regions *biomartR* package for R was used. Differences in ROHi and RHZ between populations were used as genetic distances as a source to build a rooted dendrogram by using optimal leaf ordering (OLO) for hierarchical clustering available in the *heatmaply* R package^44^. The OLO clusters similar groups (or leaves) taken from the UPGMA (Unweighted Pair Grouping with Arithmetic Mean) algorithm and yields the leaf order that maximizes the sum of the similarities of adjacent leaves in the ordering^45^.

### Islands of Runs of Homozygosity (ROHi)

ROHi are defined as regions in the genome where the proportion of individuals of a population have ROH in a specific region that is more than expected by a binomial distribution. In order to search for ROHi a sliding window of 100 Kb was used. In every 100 Kb genomic window the number of people with ROH was obtained; and to know if a specific genomic window has a significant enrichment of ROH across the population, a binomial test with P < 2×10^−7^ with Bonferroni correction for 2500 windows was applied. According to this procedure two variables could introduce bias when comparing populations across the globe: different population sizes and ROH background. In order to mitigate this source of bias the following steps were followed. Firstly, ROH of all the populations by geographical area and admixture were collapsed creating the following groups: Europe (n=474 individuals), Eastern Asia (n=459 individuals), Admixed African-American (n=121 individuals), Western Africa (n=319 individuals), Africa Guinea Gulf (n=299 individuals), Horn of Africa (n=107 individuals), Eastern Africa (n=570 individuals), Southern Africa (n=217 individuals), Khoe and San (n=148 individuals) and Admixed Hispanic-American (n=184 individuals). Secondly, ROH from 100 people in each group were resampled 100 times. Thirdly, statistically significant windows were obtained following the above methodology. Finally, consecutive windows found to be statistically significant in at least 50 resampling events were considered as part of the same ROHi.

In order to compare ROHi between populations it was considered that two ROHi from two different populations are indeed the same ROHi if they share at least 50% of their length. Results were compared using an alternative value of 75% without significant changes (data not shown).

### Regions of Heterozygosity (RHZ)

RHZ are defined as regions in the genome where < 5% of individuals in a population have ROH. In order to search for RHZ an extra step of QC consisting of removing the SNPs in LD using PLINK was performed before calling for ROH. For this analysis, ROH longer than 100 Kb were called using 25 SNPs per window in PLINK. With this procedure all ROH longer than 100 Kb, independent of their origin (LD or IBD), were detected with accuracy due to the SNP coverage available. Removing SNPs in LD, on average 1.1M SNPs were still available for every population, enabling detection of ROH longer than 100 Kb (2.8 Kb per SNP, in 100 Kb would be on average 35 SNPs, and a window of 25 SNPs is appropriate to cover genomic regions with less than the average number of SNPs). Once every ROH is called, it is straightforward to obtain regions outside ROH, and since SNPs in LD were pruned, these regions will be mostly heterozygous. In order to only identify informative heterozygous haplotypes, regions that have anomalous, unstructured, high signal/read counts in next generation sequence experiments were removed. These 226 regions, called ultra-high signal artifact regions, include high map-ability islands, low map-ability islands, satellite repeats, centromere regions, snRNA and telomeric regions^46^. Regions not covered by the Human Omni Chip 2.5 were also removed from the analyses (Like p arms of chromosomes 13, 14, 15, 21 and 22). By moving a 100 Kb window through the genome, two different cutoffs were considered to call RHZ in each window: no individual is in homozygosis (RHZ 0%) or 5% or less of the individuals are in homozygosis (RHZ 5%). Consecutive windows that fulfill this requirement were considered part of the same RHZ.

## Results

### Comparison of different ROH sizes across world populations

Data analysis of mean total lengths (sum of ROH) of different ROH length classes were plotted (Figure 2). Three different situations were considered: ROH<1Mb, 1<ROH<4Mb and ROH>4Mb. Within Sub-Saharan Africa (SSA), Figure 2A shows different scenarios for short (<1.Mb) and long (>4Mb) ROH: short ROH, unlike the long ROH, display differences between regions and commonality among then. The populations with the longest average sum of short ROH are from the Horn of Africa (Amhara, Oromo, Somali). Populations from Western Africa, Gulf of Guinea, Eastern Africa and Southern Africa, in this order and with slight differences, have intermediate levels of short ROH, and Colored populations from South Africa are the ones with the lowest levels of short ROH. Populations from these regions are reasonably homogeneous, unlike the Khoe and San populations. A completely different situation arises when long ROH (> 4 Mb) are considered, in this case no population or geographic structure is observed. Three populations, Wolof and Fula, from western Africa, and Somali from the Horn of Africa, present the largest mean total length. Differences between long and short ROH can also been seen when considering populations around the world (Figure 2B). African populations have the smallest mean total length of ROH, but this applies only to short ROH. When considering long ROH, African populations like the Wolof, Fula and Somali have mean total lengths larger than most of the KGP populations. Just the indigenous but partially admixed populations from Lima, Peru (PEL), had a larger mean total ROH length. Interestingly, for the vast majority of the populations the mean total length of very short ROH (0.3 to 0.5 Mb) is several times larger than the mean total length for long ROH (> 4Mb). This is not the case for the Khwe, Wolof and Fula populations.

**Figure 2.**
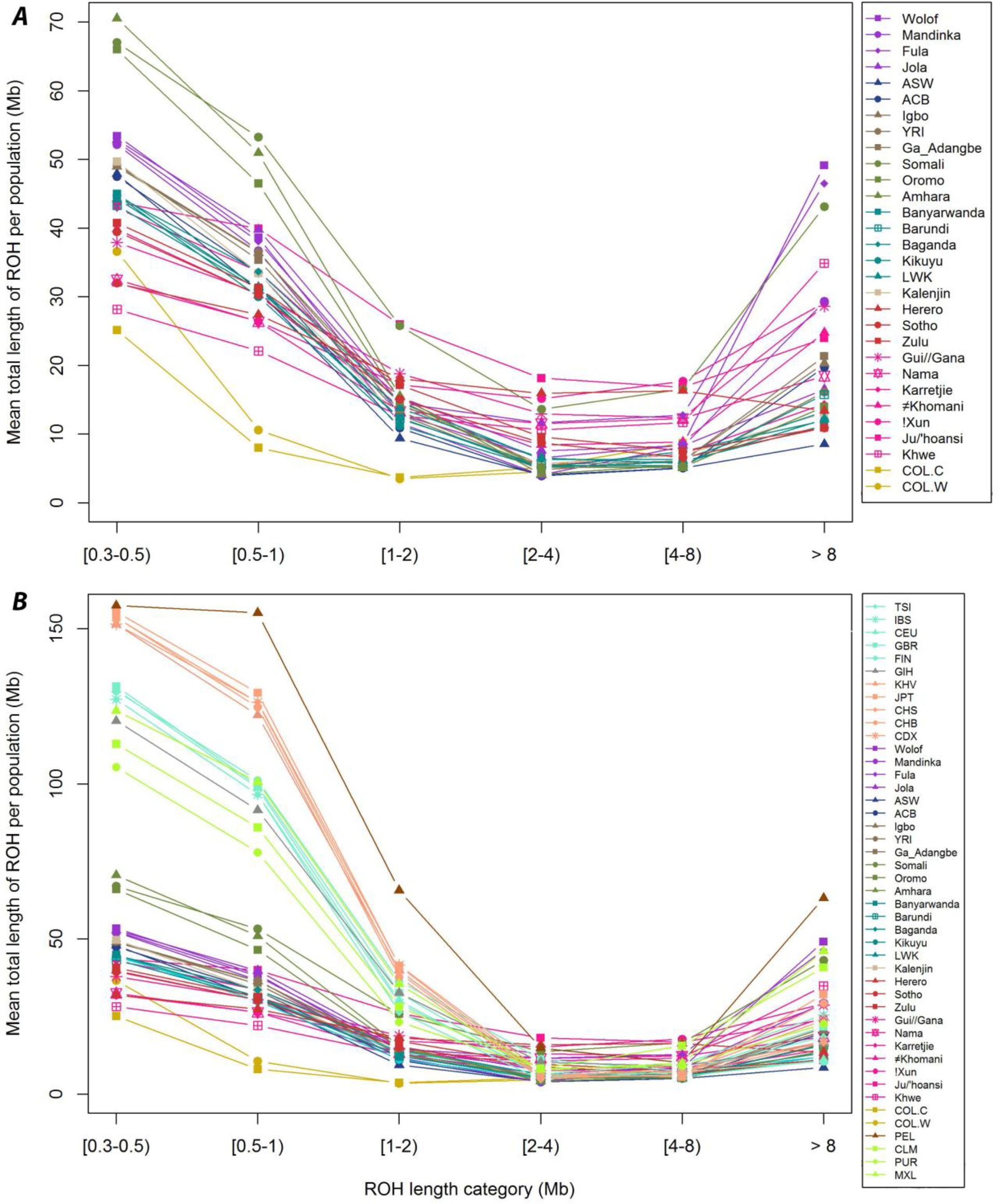
Mean total length of ROH over 6 classes of ROH tract lengths. ROH classes: 0.3≤ ROH<0.5 Mb, 0.5≤ROH<1 Mb, 1≤ROH<2 Mb, 2≤ROH<4 Mb, 4≤ROH<8 Mb and ROH≥8Mb. A. Sub-Saharan African populations and admixture populations with African ancestry (ASW and ACB, shown in dark blue). Color coding corresponds to the legend in Figure 1. B. All populations from the KGP, AGVP and Schlesbusch et al. 2012. European populations are shown in aquamarine, Southern Asian population (GIH) is shown in grey, Eastern Asia populations are shown in light salmon, South America population (PEL) is shown in dark orange, admixture Hispanic – American populations are shown in light green.

Medium size ROH (ROH between 1 a 4 Mb) (Figure 2) also reveals interesting differences. At a population level, the Khoe and San groups like Ju/’hoansi, !Xun and Khwe, have a higher mean total length for ROH from 2 to 8 Mb, even higher than PEL. Medium size ROH also show an interesting global pattern: a considerable reduction in mean total length of ROH can be seen for all populations across the globe, and there are no big differences between populations for mean total length for those ROH length classes. Considering the limitations of the KGP dataset to represent world populations, the HGDP was added to the exploratory analysis (Figure 3). In this dataset it is possible to find very isolated populations from Oceania and America and a better representation of Asian populations. Figure 3 shows the same tendency even in very isolated populations, like the African Hadza, who also have a reduction in medium size ROH.

**Figure 3.**
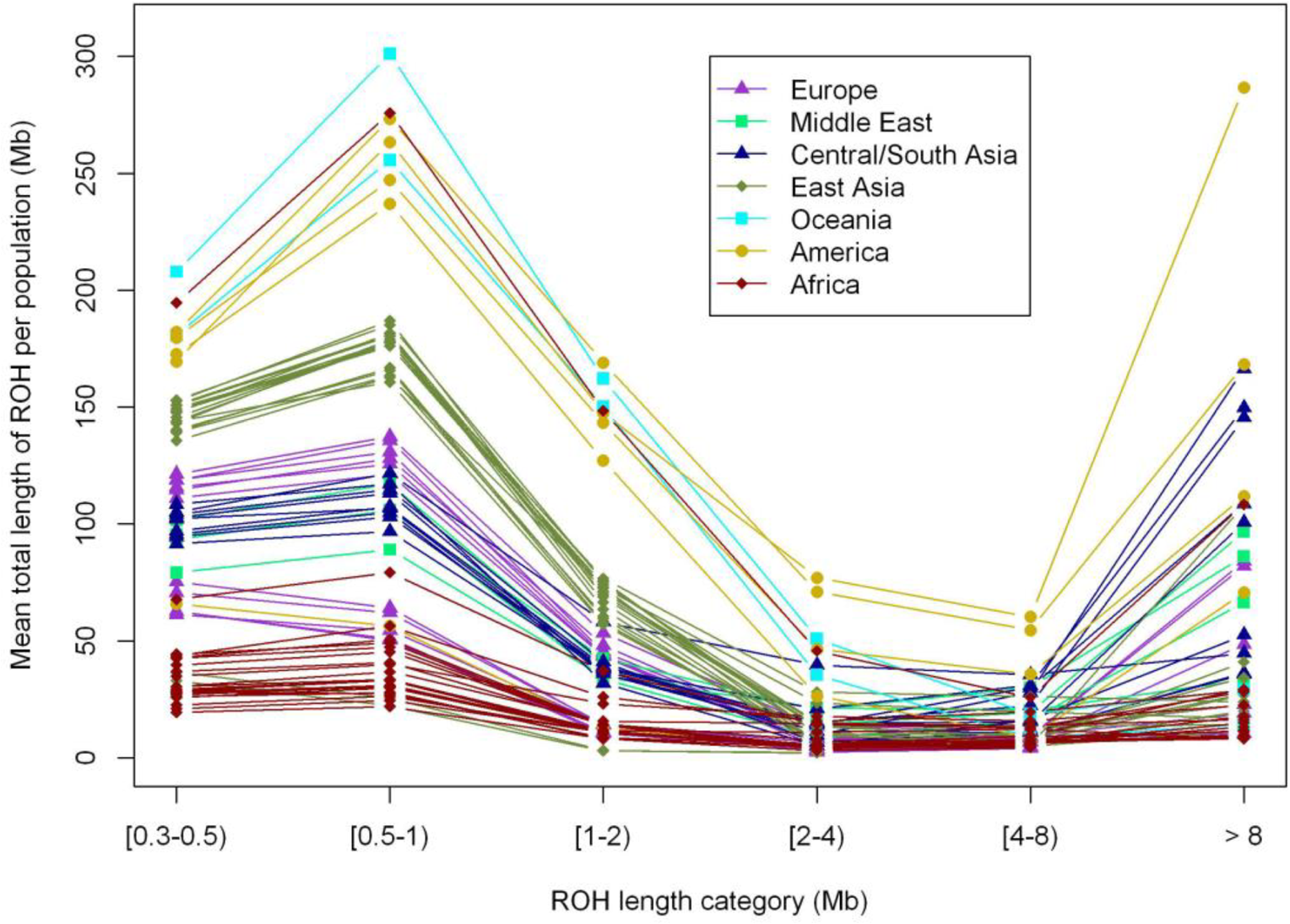
Mean total length of ROH over 6 classes of ROH tract lengths for the merged dataset of AGVP, KGP, Schlesbusch and HGDP (worldata.03, see text in the Materials and Methods section). Europe populations are shown in deep purple, Middle east populations are shown in deep purple, Middle east populations are shown in light green, Central and South Asia populations are shown in dark blue, Eastern Asia population are shown in dark green, Oceanic populations are shown in light blue, American populations are shown in yellow and African ones are shown in red.

### Violin Plots: Exploratory data analysis and non-parametrical comparisons

Using violin plots, it is possible to examine the distribution of ROH in SSA. Figure 4 represents the distributions and medians, complemented with the mean and standard deviations in Table 1. Within SSA the population with the greatest number of ROH (for ROH longer than 1.5Mb) is the Khoe-San Ju/’hoansi (median=14.5, mean=15.1). Considering populations from around the globe, only PEL has a higher number of ROH (median=18, mean=17.9). The Khoe-San populations, in general, are the ones with a higher number of ROH in SSA; however, they also show great variability. For example, both San Tuu speaker populations, ≠Khonami and Karretjie, have a considerably smaller number of ROH (median=6, mean=6.7 and median=4, mean=5.15 respectively). Besides Khoe and San populations we observe other populations like Somali and Herero with a large number of ROH (median=13, mean=13.6 and median=13.5, mean=13.3 respectively). Among SSA we find great variability, for example, populations like the Fula have a smaller number of ROH (median=6, mean=8.4) but with a long right tail (sd=7.2) which indicates great variability within the population (Figure 4). These right tails of the distribution are even longer when considering the mean size of ROH (ROH>1.5Mb). Populations from Western Africa (Fula, Wolof and Mandinka) present the longest right tails along with the TSI population from the Iberia peninsula in Europe (Table1).

**Table 1.**
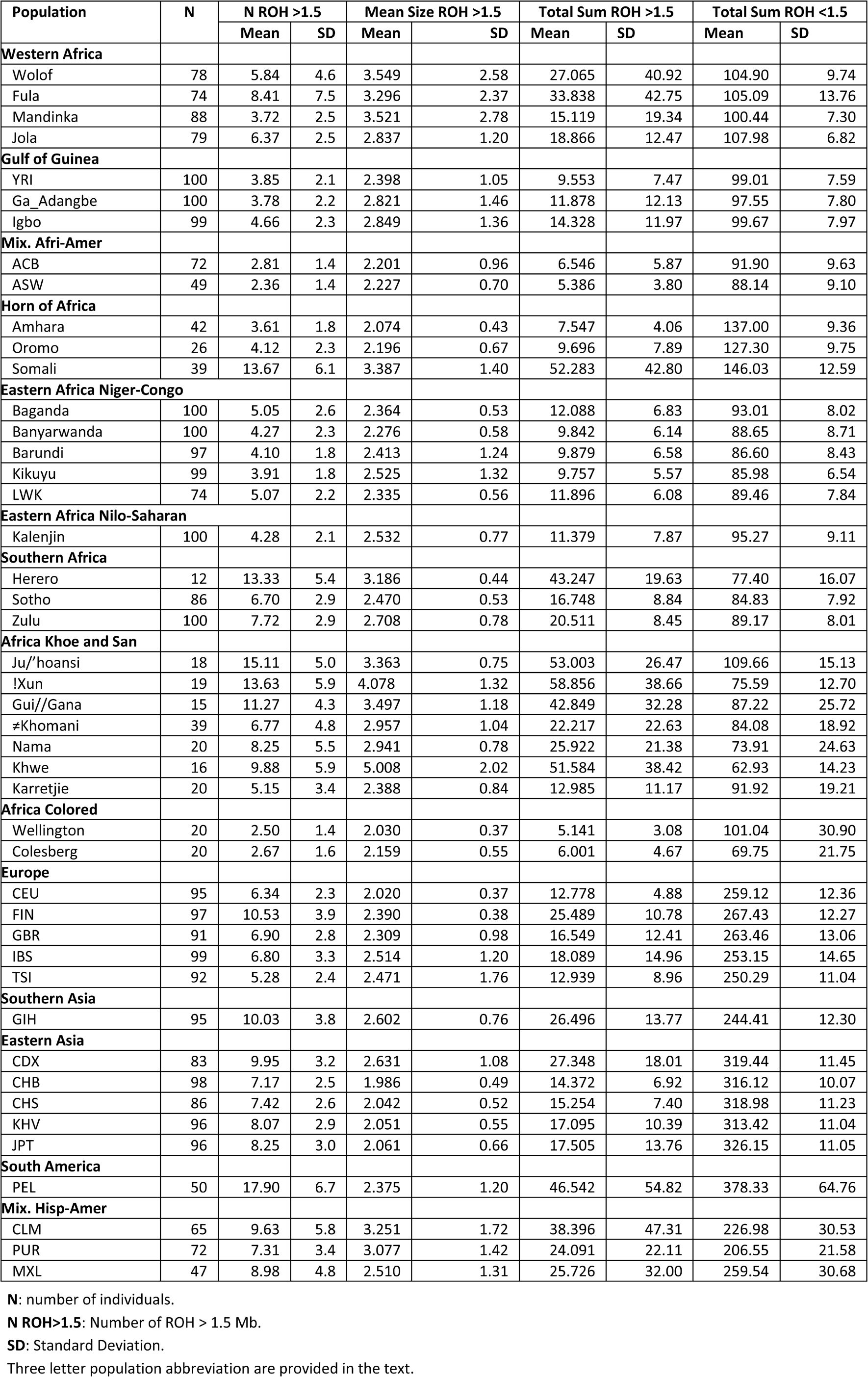
*Number, size distribution and sum of ROH (above and below 1.5Mb) across global regions and according to population.*

| Population | N | N ROH >1.5 |  | Mean Size ROH >1.5 |  | Total Sum ROH >1.5 |  | Total Sum ROH <1.5 |  |
| --- | --- | --- | --- | --- | --- | --- | --- | --- | --- |
|  |  | Mean | SD | Mean | SD | Mean | SD | Mean | SD |
| Western Africa |  |  |  |  |  |  |  |  |  |
| Wolof | 78 | 5.84 | 4.6 | 3.549 | 2.58 | 27.065 | 40.92 | 104.90 | 9.74 |
| Fula | 74 | 8.41 | 7.5 | 3.296 | 2.37 | 33.838 | 42.75 | 105.09 | 13.76 |
| Mandinka | 88 | 3.72 | 2.5 | 3.521 | 2.78 | 15.119 | 19.34 | 100.44 | 7.30 |
| Jola | 79 | 6.37 | 2.5 | 2.837 | 1.20 | 18.866 | 12.47 | 107.98 | 6.82 |
| Gulf of Guinea |  |  |  |  |  |  |  |  |  |
| YRI | 100 | 3.85 | 2.1 | 2.398 | 1.05 | 9.553 | 7.47 | 99.01 | 7.59 |
| Ga_Adangbe | 100 | 3.78 | 2.2 | 2.821 | 1.46 | 11.878 | 12.13 | 97.55 | 7.80 |
| Igbo | 99 | 4.66 | 2.3 | 2.849 | 1.36 | 14.328 | 11.97 | 99.67 | 7.97 |
| Mix. Afri-Amer |  |  |  |  |  |  |  |  |  |
| ACB | 72 | 2.81 | 1.4 | 2.201 | 0.96 | 6.546 | 5.87 | 91.90 | 9.63 |
| ASW | 49 | 2.36 | 1.4 | 2.227 | 0.70 | 5.386 | 3.80 | 88.14 | 9.10 |
| Horn of Africa |  |  |  |  |  |  |  |  |  |
| Amhara | 42 | 3.61 | 1.8 | 2.074 | 0.43 | 7.547 | 4.06 | 137.00 | 9.36 |
| Oromo | 26 | 4.12 | 2.3 | 2.196 | 0.67 | 9.696 | 7.89 | 127.30 | 9.75 |
| Somali | 39 | 13.67 | 6.1 | 3.387 | 1.40 | 52.283 | 42.80 | 146.03 | 12.59 |
| Eastern Africa Niger-Congo |  |  |  |  |  |  |  |  |  |
| Baganda | 100 | 5.05 | 2.6 | 2.364 | 0.53 | 12.088 | 6.83 | 93.01 | 8.02 |
| Banyarwanda | 100 | 4.27 | 2.3 | 2.276 | 0.58 | 9.842 | 6.14 | 88.65 | 8.71 |
| Barundi | 97 | 4.10 | 1.8 | 2.413 | 1.24 | 9.879 | 6.58 | 86.60 | 8.43 |
| Kikuyu | 99 | 3.91 | 1.8 | 2.525 | 1.32 | 9.757 | 5.57 | 85.98 | 6.54 |
| LWK | 74 | 5.07 | 2.2 | 2.335 | 0.56 | 11.896 | 6.08 | 89.46 | 7.84 |
| Eastern Africa Nilo-Saharan |  |  |  |  |  |  |  |  |  |
| Kalenjin | 100 | 4.28 | 2.1 | 2.532 | 0.77 | 11.379 | 7.87 | 95.27 | 9.11 |
| Southern Africa |  |  |  |  |  |  |  |  |  |
| Herero | 12 | 13.33 | 5.4 | 3.186 | 0.44 | 43.247 | 19.63 | 77.40 | 16.07 |
| Sotho | 86 | 6.70 | 2.9 | 2.470 | 0.53 | 16.748 | 8.84 | 84.83 | 7.92 |
| Zulu | 100 | 7.72 | 2.9 | 2.708 | 0.78 | 20.511 | 8.45 | 89.17 | 8.01 |
| Africa Khoe and San |  |  |  |  |  |  |  |  |  |
| Ju/'hoansi | 18 | 15.11 | 5.0 | 3.363 | 0.75 | 53.003 | 26.47 | 109.66 | 15.13 |
| !Xun | 19 | 13.63 | 5.9 | 4.078 | 1.32 | 58.856 | 38.66 | 75.59 | 12.70 |
| Gui//Gana | 15 | 11.27 | 4.3 | 3.497 | 1.18 | 42.849 | 32.28 | 87.22 | 25.72 |
| ≠Khomani | 39 | 6.77 | 4.8 | 2.957 | 1.04 | 22.217 | 22.63 | 84.08 | 18.92 |
| Nama | 20 | 8.25 | 5.5 | 2.941 | 0.78 | 25.922 | 21.38 | 73.91 | 24.63 |
| Khwe | 16 | 9.88 | 5.9 | 5.008 | 2.02 | 51.584 | 38.42 | 62.93 | 14.23 |
| Karretjie | 20 | 5.15 | 3.4 | 2.388 | 0.84 | 12.985 | 11.17 | 91.92 | 19.21 |
| Africa Colored |  |  |  |  |  |  |  |  |  |
| Wellington | 20 | 2.50 | 1.4 | 2.030 | 0.37 | 5.141 | 3.08 | 101.04 | 30.90 |
| Colesberg | 20 | 2.67 | 1.6 | 2.159 | 0.55 | 6.001 | 4.67 | 69.75 | 21.75 |
| Europe |  |  |  |  |  |  |  |  |  |
| CEU | 95 | 6.34 | 2.3 | 2.020 | 0.37 | 12.778 | 4.88 | 259.12 | 12.36 |
| FIN | 97 | 10.53 | 3.9 | 2.390 | 0.38 | 25.489 | 10.78 | 267.43 | 12.27 |
| GBR | 91 | 6.90 | 2.8 | 2.309 | 0.98 | 16.549 | 12.41 | 263.46 | 13.06 |
| IBS | 99 | 6.80 | 3.3 | 2.514 | 1.20 | 18.089 | 14.96 | 253.15 | 14.65 |
| TSI | 92 | 5.28 | 2.4 | 2.471 | 1.76 | 12.939 | 8.96 | 250.29 | 11.04 |
| Southern Asia |  |  |  |  |  |  |  |  |  |
| GIH | 95 | 10.03 | 3.8 | 2.602 | 0.76 | 26.496 | 13.77 | 244.41 | 12.30 |
| Eastern Asia |  |  |  |  |  |  |  |  |  |
| CDX | 83 | 9.95 | 3.2 | 2.631 | 1.08 | 27.348 | 18.01 | 319.44 | 11.45 |
| CHB | 98 | 7.17 | 2.5 | 1.986 | 0.49 | 14.372 | 6.92 | 316.12 | 10.07 |
| CHS | 86 | 7.42 | 2.6 | 2.042 | 0.52 | 15.254 | 7.40 | 318.98 | 11.23 |
| KHV | 96 | 8.07 | 2.9 | 2.051 | 0.55 | 17.095 | 10.39 | 313.42 | 11.04 |
| JPT | 96 | 8.25 | 3.0 | 2.061 | 0.66 | 17.505 | 13.76 | 326.15 | 11.05 |
| South America |  |  |  |  |  |  |  |  |  |
| PEL | 50 | 17.90 | 6.7 | 2.375 | 1.20 | 46.542 | 54.82 | 378.33 | 64.76 |
| Mix. Hisp-Amer |  |  |  |  |  |  |  |  |  |
| CLM | 65 | 9.63 | 5.8 | 3.251 | 1.72 | 38.396 | 47.31 | 226.98 | 30.53 |
| PUR | 72 | 7.31 | 3.4 | 3.077 | 1.42 | 24.091 | 22.11 | 206.55 | 21.58 |
| MXL | 47 | 8.98 | 4.8 | 2.510 | 1.31 | 25.726 | 32.00 | 259.54 | 30.68 |

N: number of individuals.
N ROH>1.5: Number of ROH > 1.5 Mb.
SD: Standard Deviation.
Three letter population abbreviation are provided in the text.

**Figure 4.**
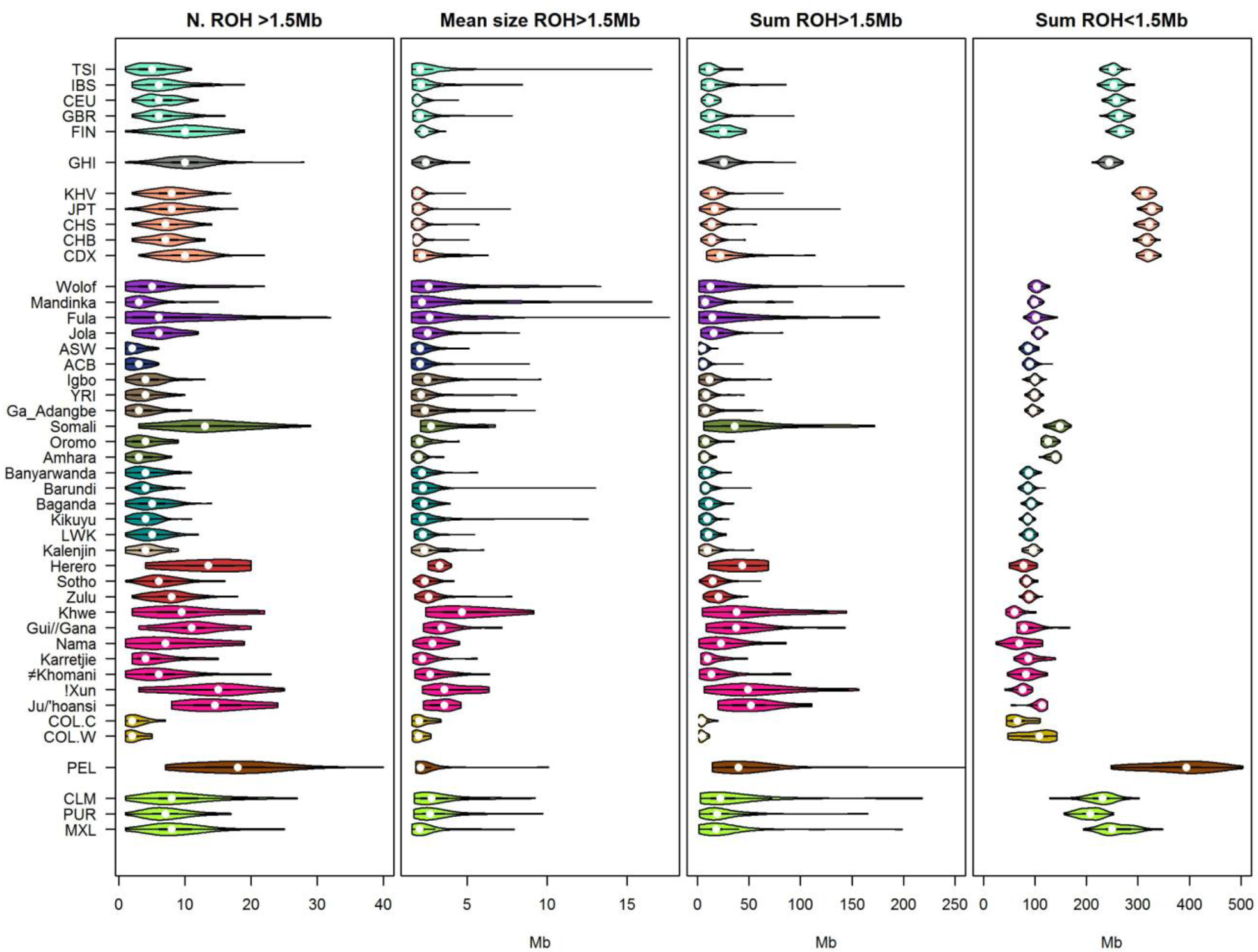
Violin plots showing the distribution of ROH within populations for the mean number of ROH longer than 1.5Mb, mean size of ROH longer than 1.5Mb, mean total sum of ROH longer than 1.5Mb and mean total length of ROH shorter than 1.5M. The colors are coded according to the legends of Figures 1 and 2.

Differences between the short and long ROH seen in Figure 2 are represented more clearly in Figure 4. Geographic classification and stratification can be seen for mean sum of ROH <1.5Mb: SSA populations have the lowest medians (Figure S8), and within the continent, populations from the Horn of Africa have a significant higher sum of ROH as shown in Figure S7. Figure 4 and Table 1 show that, without considering Horn of Africa populations, there are no real differences between Khoe-San and the rest of the SSA populations. In Table 1 populations like the Ju/’hoansi, with a mean total sum of ROH <1.5Mb (109.66 Mb), are slightly higher than populations from Western Africa, or populations like the !Xun, Nama or Khwe with the smallest mean total sum of ROH<1.5Mb in all SSA (75.5, 73.9 and 62.9 Mb respectively) besides the Colesberg Colored population with 69.7Mb. The shapes of the violin plots for sum of ROH <1.5Mb provide additional information. In general, populations are homogeneous, with very short tails and an almost normal distribution; however, Khoe and San, Colored and populations from America present more variability. Distribution shapes are completely different for the sum of ROH >1.5Mb. When considering these ROH we observe greater variability of the distribution shapes across populations within and outside SSA. Wolof (median=12.5Mb, mean=27.1Mb, sd=40.9Mb), Fula (median=14.4Mb, mean=33.8Mb, sd=42.7 Mb) and Somali (median=35.8Mb, mean=52.3Mb, sd=42.1Mb) show especially long right tails, and just two populations outside SSA: PEL (median=39.6Mb, mean=46.5Mb, sd=54.8Mb) and CLM (median=22.2Mb, mean=38.4Mb, sd=47.3Mb) have longer tails. Khoe-San populations form a heterogeneous group, but also show long tails and widely spread distributions, indeed two populations with the highest total sum of ROH are Khoe-San: the !Xun population from Angola (median=48.9Mb, mean=58.8Mb, sd=38.7Mb) and the Ju/’hoansi from Namibia (median=51.8Mb, mean=53.0Mb, sd=109.7Mb). Figures S7 and S8 show non-parametrical pairwise statistical comparisons between SSA populations and world regions.

### Inbreeding Coefficient from ROH: F_ROH_

The genomic inbreeding coefficient from ROH was obtained as the total sum of ROH longer than 1.5Mb divided by the total length of the autosomal genome. For practical reasons a cut-off point of F_ROH_ = 0.0156 (corresponding to the mean kinship of a second cousin marriage) was set to differentiate between inbred and non-inbred individuals. In the demographic literature consanguineous marriage is usually defined as a union between individuals who are related as second cousin or closer. This arbitrary limit is based on the perception that an inbreeding coefficient below 0.0156 has biological effects not very different from those found in the general population ^47^.

Table 2 shows the mean F_ROH_, the max F_ROH_, the number and proportion (in %) of individuals with an F_ROH_ between second (F=0.0156) and first cousin (F=0.0625), and the number of individuals with an F_ROH_ higher than first cousin per population. The highest average F_ROH_ for all populations can be found in the Khoe-San, !Xun and Ju/’hoansi people with an average F_ROH_ of 0.0204 and 0.0184 respectively showing them to be the most inbred populations. Besides these two, Somali people from the Horn of Africa, the Khwe Khoe and San, the PEL population and the Gui//Gana Khoe-San (average F_ROH_=0.0181; 0.0179; 0.0162 and 0.0151 respectively) have mean F_ROH_ higher than a second cousin kinship. The individual with the highest inbreeding coefficient from ROH across all populations is a Peruvian with an F_ROH_ of 0.1400 (higher than an uncle-niece or double first cousin kinship, θ=0.125). Within SSA, only the Wolof from Western Africa has individuals with inbreeding coefficients higher than a first cousin union. Figure 5 plots the number of ROH (longer than 1.5Mb) and the total sum of ROH >1.5Mb for each SSA individual, and shows in red dashed lines conservative limits for second and first cousin inbreeding coefficient. In this figure it can be seen that, regarding F_ROH_, populations across SSA have a wide range of inbreeding coefficient. In Western Africa (Figure 5A) Wolof and Fula individuals are more dispersed across the plot, with 17.9% of Wolof and 29.7% of Fula having an F_ROH_ higher than 0.015. In contrast, Mandinka and Jola, with just 8% and 7.6% of inbred individuals, present a tighter scattering. Populations from the Gulf of Guinea and African-American admixed populations shown even tighter clustering with the ACB and ASW admixed populations being the tightest. These differences can also be seen in Eastern and Horn of Africa (Figure 5B), just the Somali people show a great dispersion, 48.7% of the sample have a F_ROH_ higher than 0.015. For Southern African populations it is possible to see the dispersion of the Khoe and San populations (Figure 5C). The 73.7%, 62.5% and 60.0% individuals of the !Xun, Khwe and Gui//Gana respectively have an F_ROH_ higher than a second cousin union. These populations therefore have a large proportion of inbred individuals, even more than the partially indigenous PEL population (56%); however due to the small population sample sizes these numbers should be viewed with caution. At the opposite end of the spectrum, Colored populations have a tight distribution with very low F_ROH_.

**Table 2.**
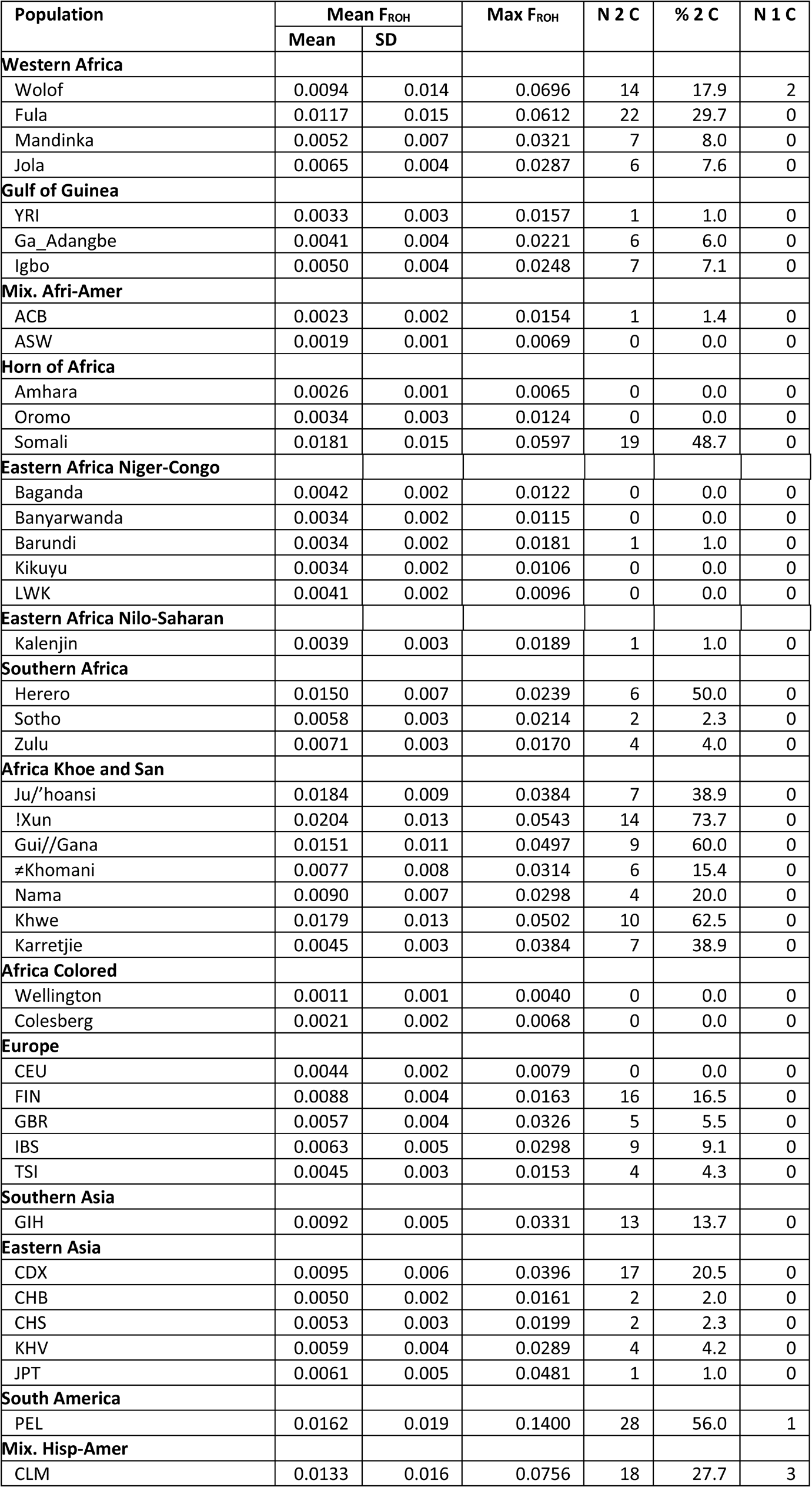
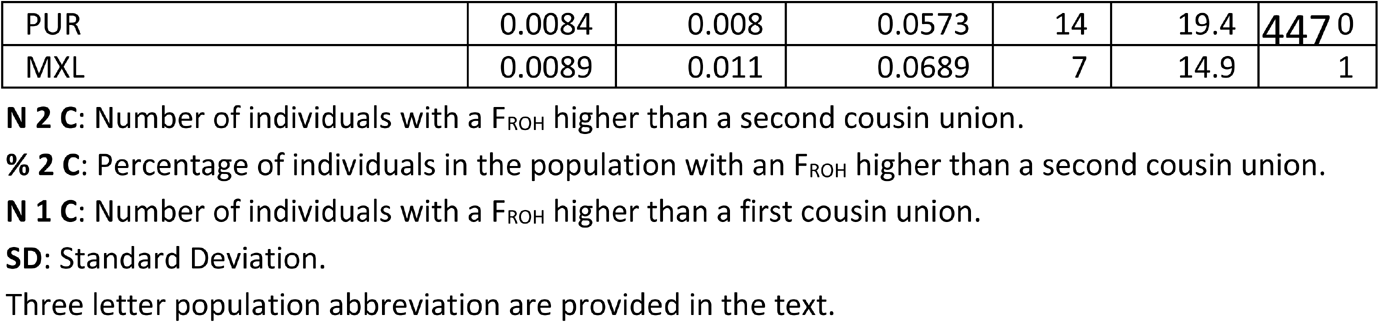
*Summary statistics for the inbreeding coefficient calculated from ROH (F_ROH_) across global regions and according to population.*

**Figure 5.**
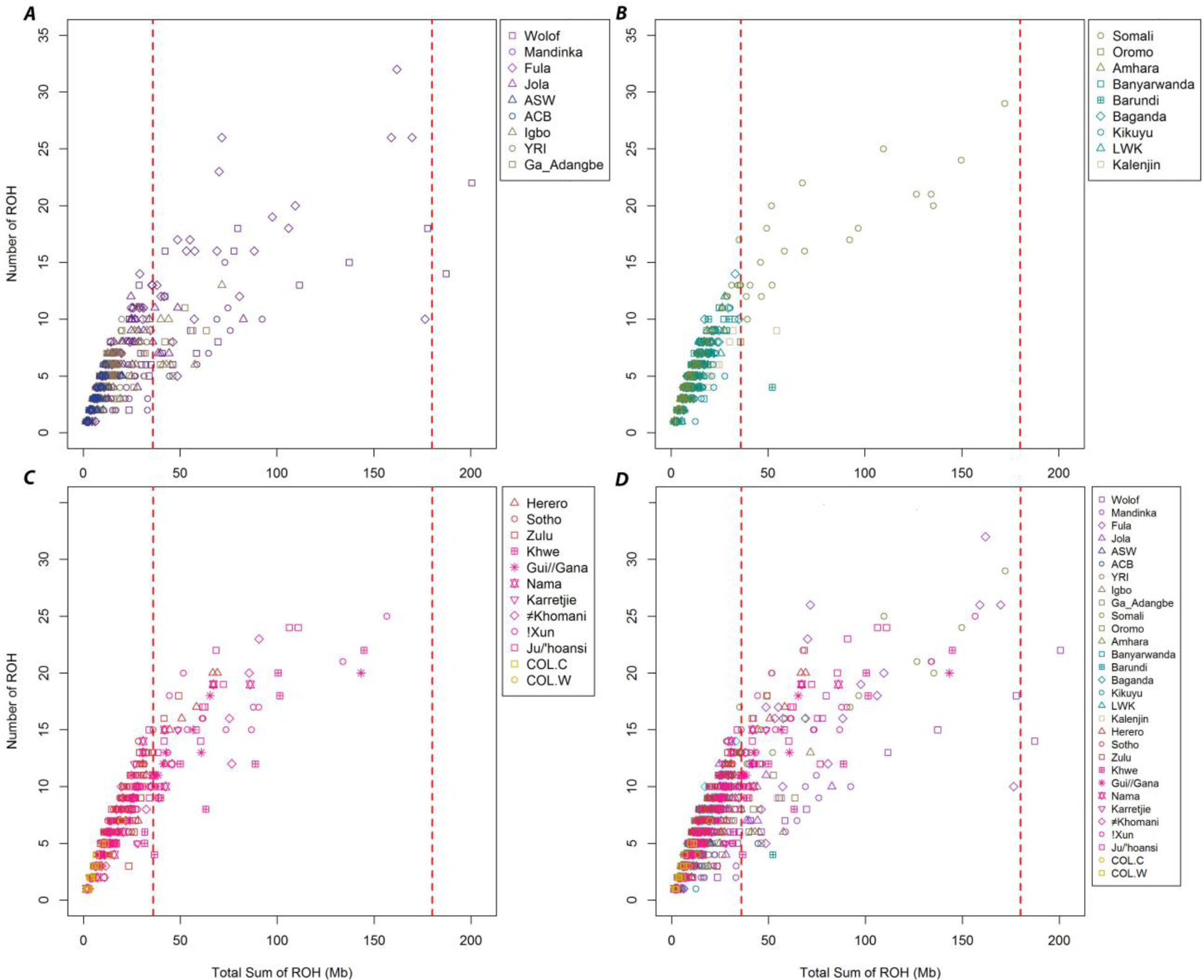
Each Sub-Sharan African individual is plotted according to their number of ROH and total sum of ROH. The perpendicular broken red lines in all the plots at X=36 and X=180, represent conservative thresholds for inbreeding coefficients of 0.0156 (second cousin offspring) and 0.0625 (first cousin offspring). A. Individuals from Western Africa and the Gulf of Guinea. B. Populations from Eastern Africa and the Horn of Africa. C. Populations from Southern Africa. D. All populations together. For color legend see figure 1 (as above)

### Discriminating between different sources of autozygosity: understanding population demographic history

Like the inbreeding coefficient calculated from a deep pedigree, F_ROH_ denotes the total inbreeding coefficient, but it does not give information regarding how that autozygosity was generated. Was it the result of cultural practices favoring related unions, or because of a low effective population size and genetic drift?

In Figure 6A the mean number of ROH (>1.5Mb) is plotted against the mean total sum of ROH (>1.5Mb) by population. The diagonal (red dashed line) was obtained by regressing both variables of the Colored population as a non-consanguineous control group. Populations falling near this diagonal line, including most of the Europeans, Asians and Africans, carry a complement of ROH derived from their continental effective population size (N_e_). The number of ROH in these populations is driven mostly by numerous short to medium ROH sizes, but longer than 1.5Mb. Under this scenario, autozygosity provoked by genetic drift will generate a large number of ROH, but short in size. On the other hand, recent inbreeding loops will produce small numbers of very long ROH which will influence the sum of ROH much more than the total number of ROH. Populations like Somali, Khwe, !Xun and to a lesser degree Fula, Wolof or CLM, which display a right shift away from the trend line in the X-axis, suggest the practice of consanguineous unions. A different approach toward differentiating the two sources of inbreeding is shown in Figure 6B. In this figure the F_IS_ in plotted against the F_ROH_ for different populations. Three different regions can be considered in this plot delimited by the diagonal, where F_IS_=F_ROH_, and the horizontal line F_IS_=0. Populations close to the diagonal line, like the Somali, have a strong component of systematic inbreeding or F_IS_, which means that the total inbreeding coefficient, F_IT_, of this population is mainly produced by a deviation from panmixia, in other words, consanguinity. Panmictic inbreeding, caused by genetic drift will be more relevant as the population gets close to the line F_IS_=0. Low N_e_, isolation and genetic drift become very relevant when populations have negative F_IS_. Under this scenario of avoidance of consanguinity and excess of heterozygotes (expected under H-W proportions), the total inbreeding coefficient of populations like PEL, Khwe, Ju/’hoansi, !Xun or Herero will be provoked by genetic isolation and genetic drift: strong F_ST_.

**Figure 6.**
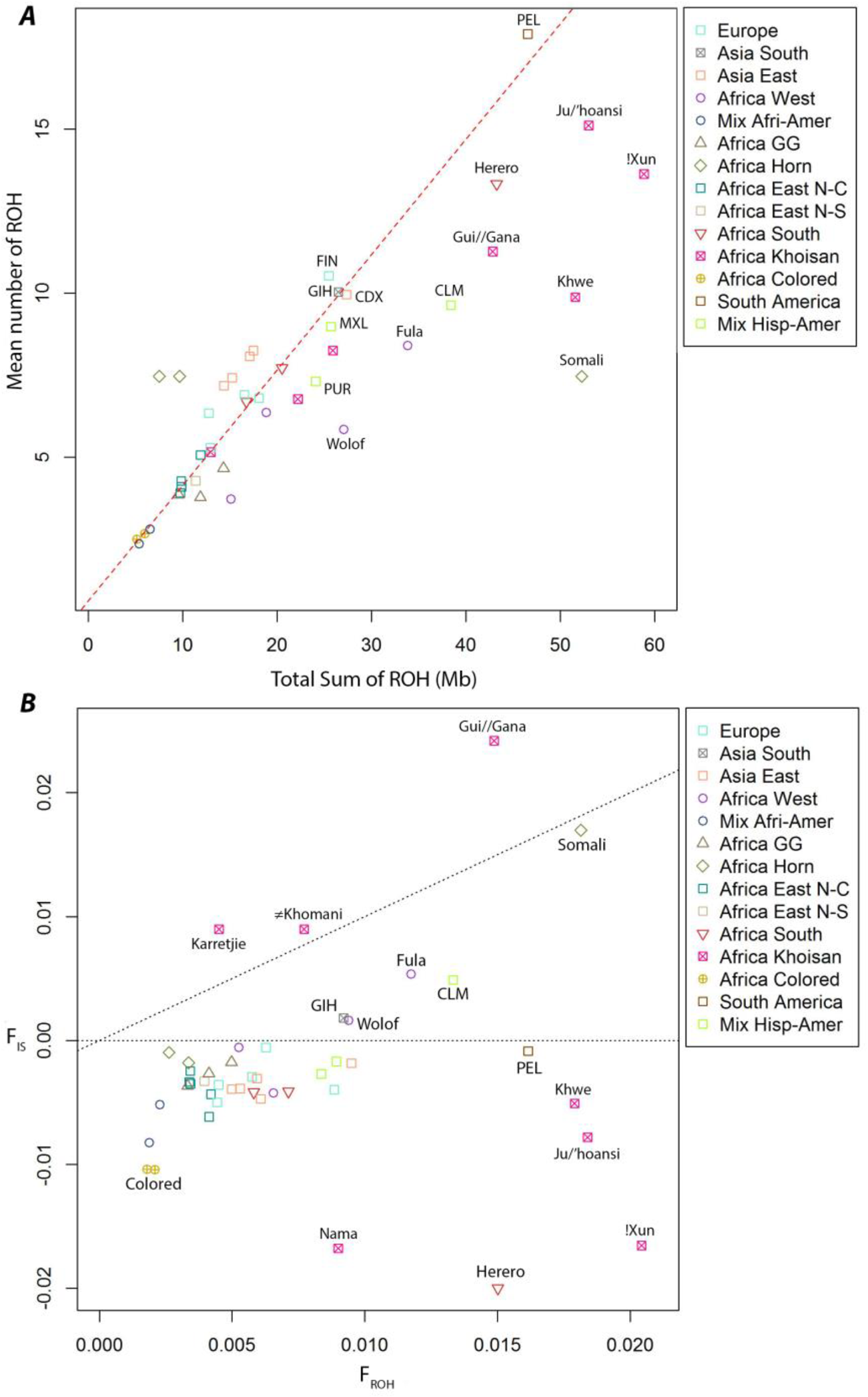
Population analysis and components of inbreeding coefficient. A. Mean number of ROH plotted versus mean total sum of ROH in Mb for the 28 populations under study (symbols according to regional groupings). Red broke line represents the regression line of the two variables (N of ROH vs Sum of ROH) for the South African Colored population (see Methods section) B. Systematic inbreeding coefficient (F_IS_) versus the inbreeding coefficient obtained from ROH (F_ROH_). Diagonal broken line represents F_IS_ = F_ROH_. Horizontal broken line represents F_IS_=0.

### Detecting the Wahlund effect

As explained above, Figure 6B has three regions: F_IS_<0, F_IS_=F_ROH_ and F_IS_>F_ROH_. Under an inbreeding context, and according to Wright F statistic, it does not make much sense for F_IS_ to be bigger than F_IT_. So, if a population presents with a larger F_IS_ other phenomena must be taken into account. Besides inbreeding, natural selection pressure and Wahlund effect can increase F_IS_; nevertheless, natural selection is an evolutionary force that can change F_IS_ locally in specific genome regions, but never at a whole genome level. The only explanation is the Wahlund effect: a deficiency of heterozygotes and excess of homozygotes provoked when subpopulations with different allele frequencies are lumped together^48^. This effect is shown in Figure 7. In this figure F_IS_ and F_ROH_ were obtained for each population and grouped by region. A perfect example is the Colored populations: when both populations are considered separately their F_IS_ is negative (-0.01 for both of them) but when combined the resulting F_IS_ is positive (+0.01). This phenomenon can be seen for the other populations and regional groups in Figure 7. When combined in their respective regional groups the resultant F_ROH_ is equal to the average of all the populations; however, the F_IS_ increases depending on the allele proportion differences between populations of a same regional group. According to this explanation, the Karretjie, ≠Khonami and Gui//Gana populations in Figure 6B may indeed be the mixture of at least two different subpopulations with different alleles frequencies.

**Figure 7.**
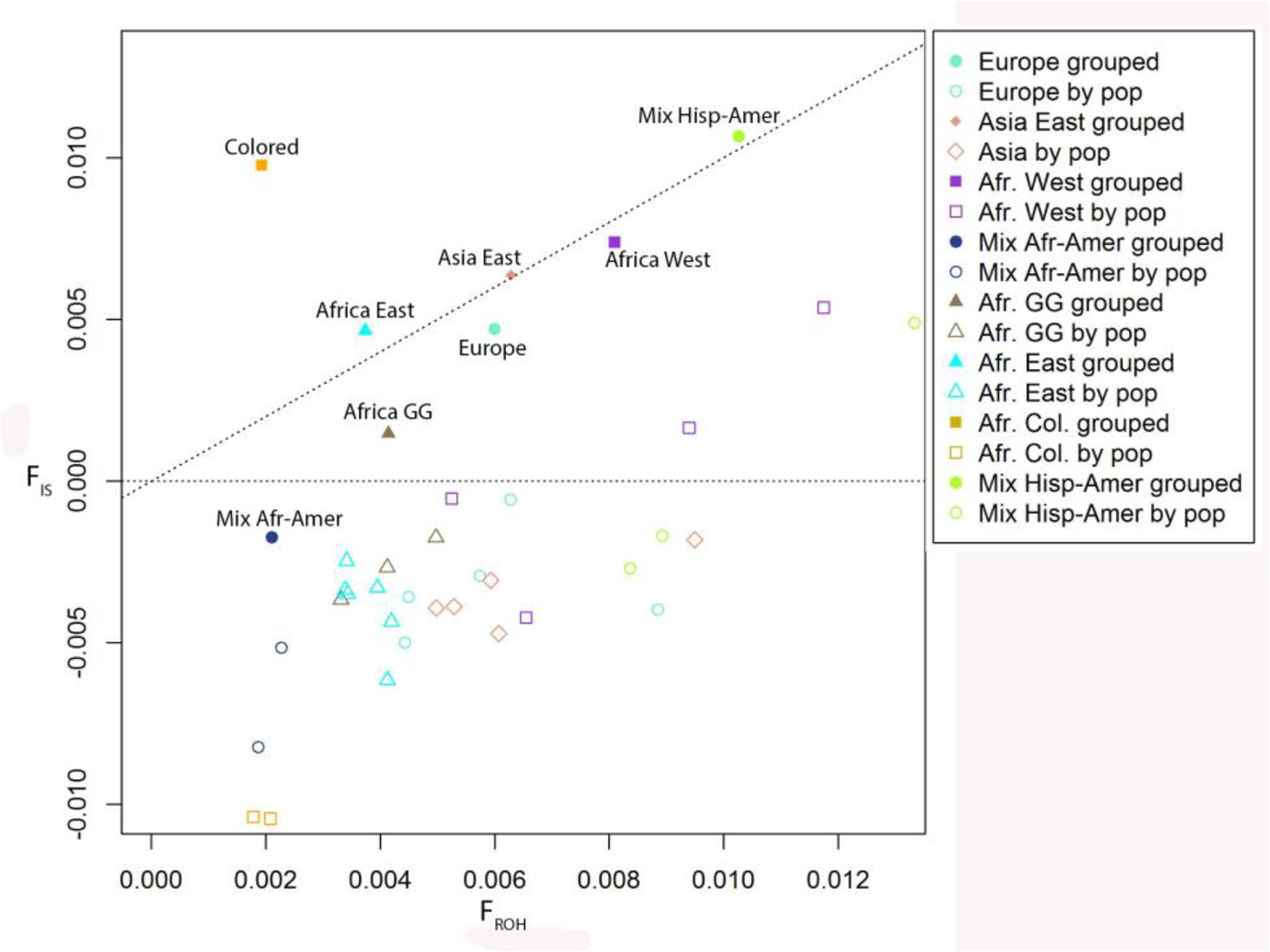
Representation of the Wahlund effect. F_IS_ and F_ROH_ values for the South African Colored population, Easter Africans, Wester Africans, Gulf of Guinea populations, mixed African-Americans, Europeans, Eastern Asia and mixed Hispanic-Americans were plotted (empty shapes). Mean F_IS_ and mean F_ROH_ per regional group are plotted and shown as solid shapes.

### Genomic distribution of Runs of Homozygosity

ROH are not randomly distributed across the genome and there are regions with a high prevalence of ROH or complete absence^7; 19; 20^. ROH islands, genomic regions with high prevalence of ROH, or regions of heterozygosity (RHZ) are analyzed by collapsing populations into their regional groups: from SSA: West, Gulf of Guinea, East, Horn of Africa, Southern Bantu and Khoe-San. From out of SSA: Europe, Eastern Asia, Hispanic-American admixed and African-American admixed. In Figure 8, ROHi and RHZ are represented for the 22 autosomal chromosomes of the Khoe and San and European groups.

**Figure 8.**
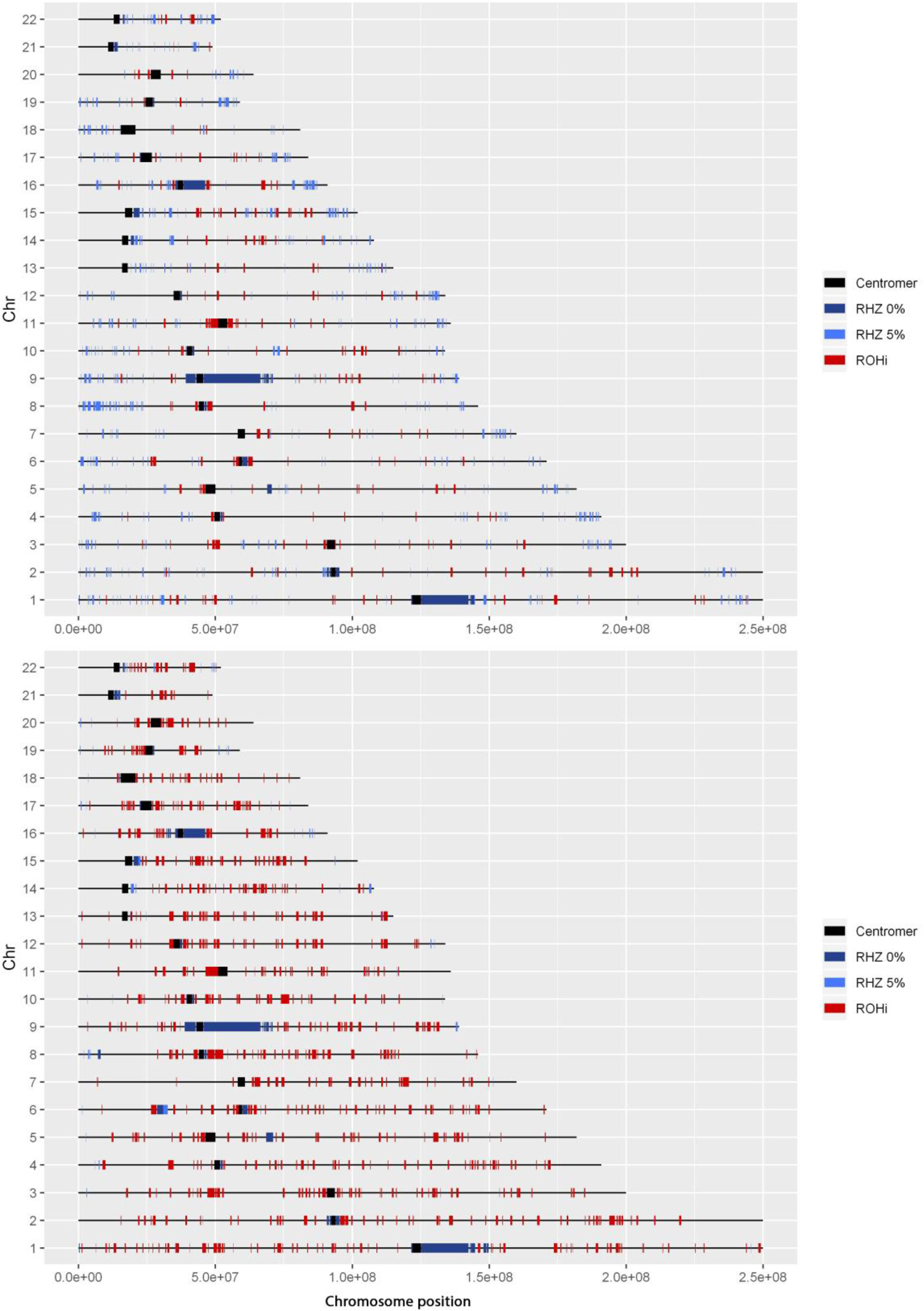
Genomic representation of the chromosomal location and size of runs of homozygosity islands (ROHi) and runs of heterozygosity (RHZ) for the Khoe and San (A) and European (B) regional groups. RHZ 0%: genomic regions where no individual in the group has a ROH. RHZ 5%: genomic regions where ≤ 5% of the population has ROH.

Within SSA, the region of the Horn of Africa has the shortest (measured in Mb and cM) but a larger number of ROHi (544) (Table 3). The Khoe and San is the group with the smallest number of ROHi, less than half (220) are of an average size. Eastern Africa has the longest ROHi measured in Mb, when measured in cM there are no big differences across SSA. Outside SSA, the Europeans form a group with the highest number of ROHi (795), 3.6 times more than the Khoe and San. Also, Europe is the group with the lager ROHi, measured in Mb and cM, with 90 ROHi larger than 1 Mb. Interestingly the African-American admixed group has almost no ROH longer than 1.5Mb, but is the group with the second highest number of ROHi. Surprisingly this group has longer ROHi with a mean size of 0.615 Mb or 0.25 cM, higher than most groups. Being the regional group from SSA with the largest number of ROHi, it seems reasonable that the Horn of Africa is the group with the least number of regions of heterozygosity, defined as regions with < 5% homozygosity (RHZ 5%). Surprisingly, this is not the case for RHZ where no individual is in homozygosity

**Table 3.** *Summary statistics for the ROH islands (ROHi) and the regions of heterozygosity (RHZ) for populations combined from different geographic regions.*

| Population | N | Number by size |  |  |  | Mean length (Mb) |  | Mean length (cM) |  | Max length |  | Mean Number of SNP |  |
| --- | --- | --- | --- | --- | --- | --- | --- | --- | --- | --- | --- | --- | --- |
|  |  | > 1.0 Mb | 1.0 - 0.5 | 0.5 - 0.3 | <0.3 Mb |  |  |  |  | Mb | cM |  |  |
|  |  |  |  |  |  | Mean | SD | Mean | SD |  |  | Mean | SD |
| Africa West |  |  |  |  |  |  |  |  |  |  |  |  |  |
| ROHi | 383 | 35 | 128 | 126 | 94 | 0.599 | 0.37 | 0.187 | 0.34 | 4.2 | 3.24 | 181.7 | 110.4 |
| RHZ 0% | 48 | 14 | 12 | 4 | 18 | 0.663 | 0.62 | 1.245 | 3.32 | 2.4 | 11.74 | 227.8 | 258.5 |
| RHZ 5% | 926 | 21 | 81 | 181 | 643 | 0.235 | 0.25 | 0.421 | 0.91 | 4.0 | 13.81 | 98.7 | 105.5 |
| Africa GG |  |  |  |  |  |  |  |  |  |  |  |  |  |
| ROHi | 370 | 40 | 117 | 138 | 75 | 0.614 | 0.41 | 0.204 | 0.40 | 4.2 | 3.77 | 184.0 | 122.5 |
| RHZ 0% | 57 | 11 | 12 | 14 | 20 | 0.691 | 0.73 | 0.742 | 1.88 | 3.6 | 7.16 | 286.8 | 301.1 |
| RHZ 5% | 1295 | 21 | 130 | 258 | 886 | 0.259 | 0.31 | 0.467 | 0.95 | 4.1 | 13.81 | 107.3 | 126.1 |
| Africa Horn |  |  |  |  |  |  |  |  |  |  |  |  |  |
| ROHi | 544 | 18 | 74 | 126 | 326 | 0.374 | 0.24 | 0.106 | 0.26 | 1.6 | 3.230 | 114.4 | 75.3 |
| RHZ 0% | 70 | 14 | 13 | 12 | 20 | 0.492 | 0.597 | 0.511 | 1.99 | 2.6 | 11.74 | 205.1 | 248.7 |
| RHZ 5% | 751 | 17 | 53 | 143 | 538 | 0.222 | 0.250 | 0.357 | 0.87 | 4 | 13.81 | 92.4 | 104.3 |
| Africa East |  |  |  |  |  |  |  |  |  |  |  |  |  |
| ROHi | 371 | 47 | 134 | 134 | 56 | 0.647 | 0.37 | 0.209 | 0.39 | 3.3 | 3.779 | 210.0 | 120.5 |
| RHZ 0% | 57 | 11 | 13 | 15 | 18 | 0.731 | 0.740 | 0.885 | 2.01 | 3.6 | 7.16 | 298.8 | 300.5 |
| RHZ 5% | 1596 | 37 | 169 | 339 | 1051 | 0.279 | 0.336 | 0.526 | 1.12 | 4.0 | 14.24 | 114.2 | 137.4 |
| Africa South |  |  |  |  |  |  |  |  |  |  |  |  |  |
| ROHi | 294 | 22 | 94 | 110 | 68 | 0.581 | 0.36 | 0.168 | 0.35 | 3.4 | 3.244 | 213.5 | 130.3 |
| RHZ 0% | 53 | 12 | 12 | 13 | 16 | 0.532 | 0.551 | 0.762 | 2.67 | 2.3 | 11.74 | 214.9 | 222.5 |
| RHZ 5% | 1300 | 32 | 152 | 342 | 774 | 0.261 | 0.261 | 0.467 | 0.95 | 2.6 | 13.81 | 105.5 | 105.3 |
| Africa Khoe and San |  |  |  |  |  |  |  |  |  |  |  |  |  |
| ROHi | 220 | 19 | 61 | 79 | 61 | 0.565 | 0.37 | 0.099 | 0.19 | 3.3 | 2.622 | 210.6 | 137.1 |
| RHZ 0% | 49 | 9 | 12 | 10 | 18 | 0.650 | 0.69 | 1.105 | 3.22 | 3.6 | 11.30 | 262.3 | 281.6 |
| RHZ 5% | 1253 | 24 | 99 | 216 | 914 | 0.237 | 0.26 | 0.387 | 0.83 | 3.7 | 11.74 | 96.0 | 103.9 |
| Mix. Af-Amer |  |  |  |  |  |  |  |  |  |  |  |  |  |
| ROHi | 689 | 72 | 221 | 252 | 144 | 0.615 | 0.41 | 0.251 | 0.34 | 5.1 | 4.233 | 146.1 | 98.6 |
| RHZ 0% | 194 | 11 | 14 | 25 | 144 | 0.294 | 0.48 | 0.270 | 1.22 | 3.6 | 13.81 | 120.9 | 196.1 |
| RHZ 5% | 1859 | 39 | 217 | 404 | 1199 | 0.284 | 0.33 | 0.511 | 1.04 | 4.6 | 14.71 | 116.5 | 134.0 |
| Europe |  |  |  |  |  |  |  |  |  |  |  |  |  |
| ROHi | 795 | 90 | 211 | 286 | 208 | 0.604 | 0.43 | 0.254 | 0.36 | 5.3 | 4.232 | 122.9 | 88.4 |
| RHZ 0% | 58 | 11 | 16 | 14 | 17 | 0.739 | 0.76 | 0.902 | 1.81 | 4.0 | 7.81 | 312.8 | 322.1 |
| RHZ 5% | 218 | 12 | 21 | 33 | 152 | 0.325 | 0.54 | 0.412 | 1.22 | 4.1 | 13.81 | 137.8 | 227.2 |
| Asia. East |  |  |  |  |  |  |  |  |  |  |  |  |  |
| ROHi | 459 | 26 | 85 | 139 | 209 | 0.466 | 0.34 | 0.128 | 0.31 | 4.1 | 3.498 | 118.8 | 87.6 |
| RHZ 0% | 57 | 11 | 15 | 14 | 17 | 0.751 | 0.78 | 1.229 | 3.07 | 4 | 11.74 | 313.5 | 328.9 |
| RHZ 5% | 195 | 14 | 16 | 33 | 132 | 0.373 | 0.62 | 0.388 | 1.13 | 4.1 | 11.75 | 155.9 | 261.1 |
| Mix. Hisp-Amer |  |  |  |  |  |  |  |  |  |  |  |  |  |
| ROHi | 645 | 56 | 171 | 205 | 213 | 0.561 | 0.40 | 0.202 | 0.34 | 5.3 | 4.232 | 144.9 | 104.6 |
| RHZ 0% | 59 | 11 | 16 | 14 | 18 | 0.726 | 0.76 | 0.846 | 1.77 | 4 | 7.16 | 302.4 | 316 |
| RHZ 5% | 273 | 12 | 22 | 48 | 191 | 0.304 | 0.49 | 0.403 | 1.10 | 4.1 | 13.81 | 126.6 | 203 |

N: number of ROHi and RHZ.
Mb: Megabases
cM: Centimorgans
SD: Standard Deviation.

The Horn of Africa actually has more of these regions than the rest of SSA groups, and only the admixed group of the African-Americans has more RHZ 0% (Table 3). Table 3 shows that for every group there are big differences between the number of RHZ 0% and 5%. These differences can be explained mainly by a drastic increase of short RHZ 5% regions (< 0.3Mb) with the outcome of a reduction in the mean length (Mb and cM) of the RHZ 5% in comparison to RHZ 0%. Table 3 also shows bigger differences between regional groups when considering RHZ in comparison to ROHi, especially in number by size and mean length. In order to appreciate differences between regional groups, three extremely long RHZ 0%, shared by all groups, were removed before constructing Table 3. These three RHZ 0% are located in Chr1 (1253+E05 to 1425+E05; 17.3Mb), Chr9 (457+E05 to 664+E05; 20.8Mb) and Chr16 (384+E05 to 463+E05; 8Mb).

Tables 4 and 5 show the positions, lengths and presence of protein coding genes for the five most common ROHi per regional group. Almost every ROHi has at least one protein coding gene, just two ROHi from the African Khoe and San and one ROHi in Hispanic-American admixed regional groups include no protein coding genes. Among the genes listed in Tables 4 and 5 there are some already described to be under positive selection pressure. Hence, there are genes related to brain development: *GPHN*^49; 50^, *PCDH17*^49^, *DARS*^49; 51^, *SCFD2*^49; 52^, *KIAA0319L*^49^, *EXOC6B*^49; 53^, *SLC30A9*^49; 53^, *CPA6*^54^, *DOCK3*^50; 55^, *CASC4*^50^ or *APBA2*^53; 56^; involved in cancer or tumor processes: *ZCCHC11*^49; 50^, *SPOCK1*^49^, *BCAS3*^49; 53^, *OLFML*^57^, *EIF2S1*^49; 57^, *MPP5*^49; 57^, *CXCR4*^51^; skin conditions: *EDAR*^49; 53; 58,^ *NOMO1*^59^; color of the eye in Europeans: *HERC2*^56^; spermatogenesis: *M1AP*, Fanconi anaemia *FANCC*^60^; pulmonary fibrosis: *PARN*^53^; congenital blindness: *TRPM1*^53^; mitochondrial disorders: *MRPS23*^49^; Charcot-Marie tooth disease: *PLEK*^49; 53^; and other metabolic and cellular processes (including *SH3RF*^49^, *CUEDC1*^49^, *GOLGA8G*^51^, *PC*^50^). Many of these ROHi with genes under positive selection are shared by more than one regional group. Without being exhaustive, the ROHi with the *FANCC* gene is present in all the SSA populations but not outside this region: 28.5% of the Western Africa population has an ROH including this gene, 29.2% of the Gulf of Guinea populations, 19.5% of the Eastern Africa regional group, 23.6% of the people from the Horn of Africa, 17.3% of the population from Southern Africa, 14.4 of the Khoe and San population and 26.9% of the admixed African-American populations. Another example shared by all SSA, except the Khoe and San populations, is the ROHi with the *GPHN* gene: 21.7% of prevalence in Western Africa, 17.8% in the Gulf of Guinea, 22.2% in Eastern Africa, 26.1% in the Africa Horn, 14.9% in Southern Africa and 20.3% of prevalence in the African-American admixed populations. ROHi with genes under positive selection were either present in all the populations like the *BCAS3* gene, or just present in only one regional group like *HERC2* or *EDAR*, in Europe and Eastern Asia respectively. Worthy of comment is the presence of an ROHi near the *LCT* gene in Europe and Eastern Africa; 38.8% and 19.9% of the European and Eastern Africa individuals have a ROHi in this gene, but not in other SSA populations.

**Table 4.**
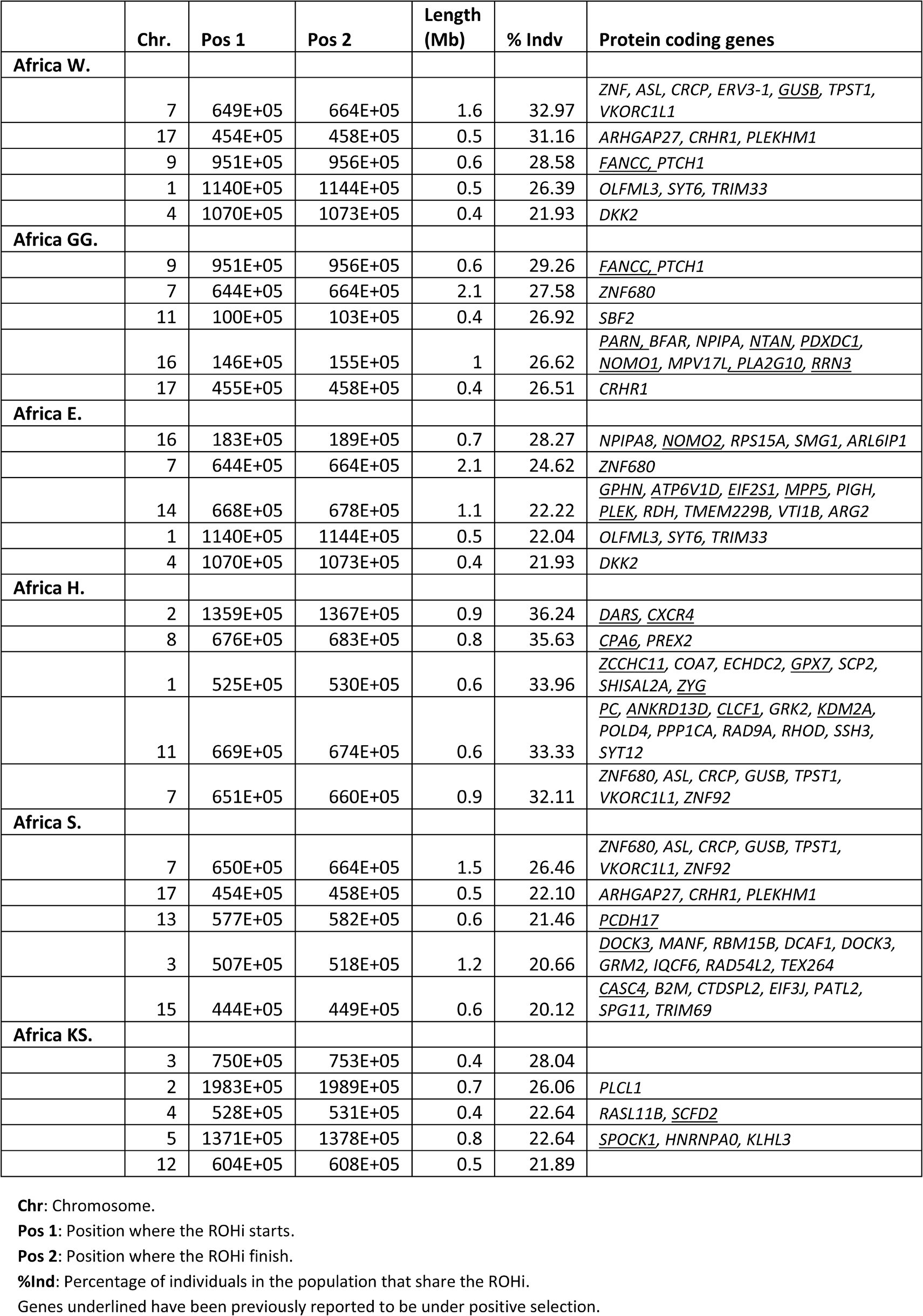
*Location, length, percentage of individuals with ROH for the ROH island and protein coding genes of the five most prevalent ROH islands in the Sub-Saharan African regional groups.*

**Table 5.**
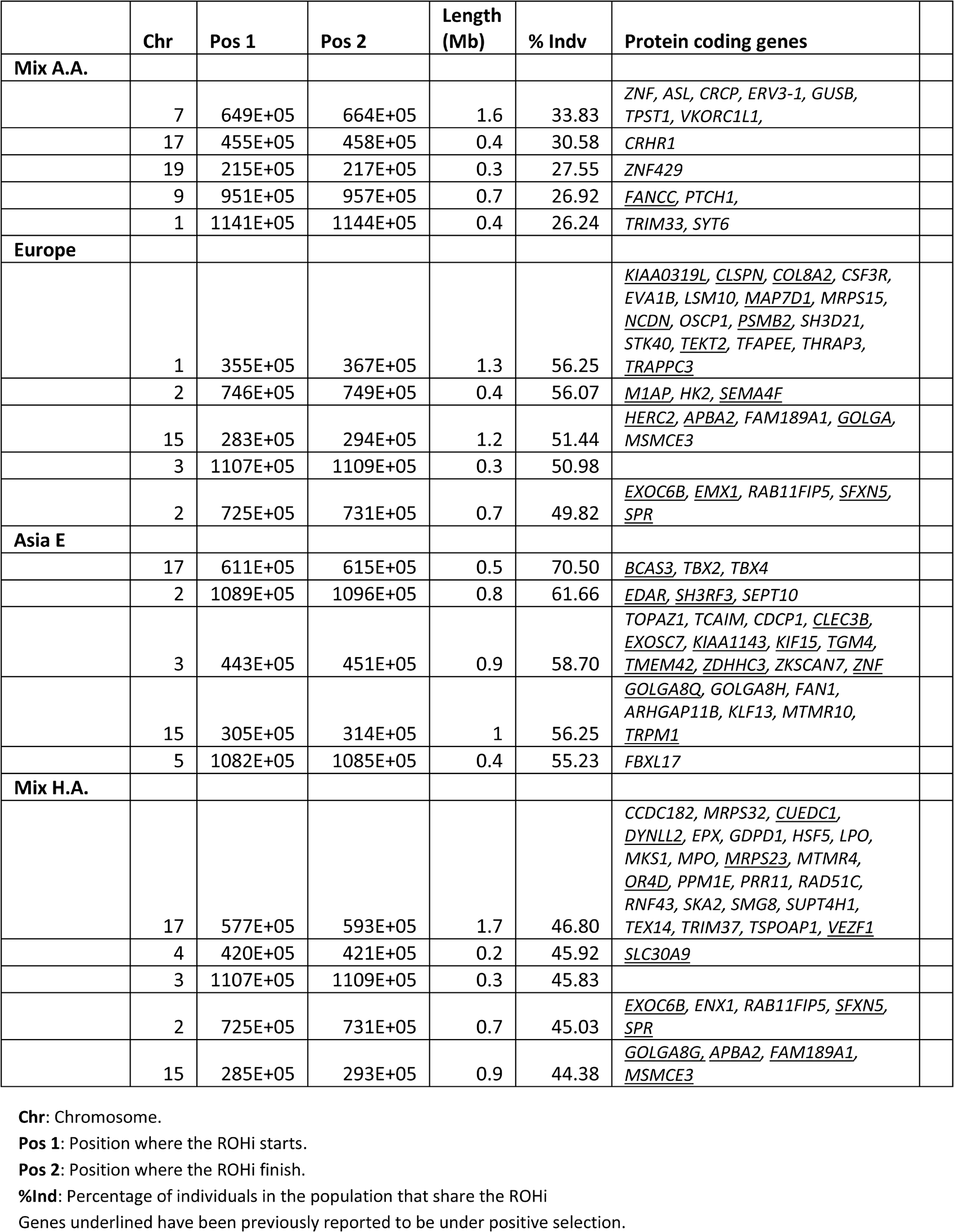
*Location, length, percentage of individuals with ROH for the ROH island and protein coding genes of the five most prevalence ROH islands in the non-African regional groups.*

Table 6 shows the three longest RHZ 5%, with the presence of protein coding genes for every regional population group. In order to build this table, the three longest RHZ 0%, present in all regional groups, were removed. These three RHZ 0% (Chr1, Chr9 and Chr16) have practically no protein coding genes, just the *SPATA31*^61^ subfamily A member 5 gene on Chr9 that is involved in spermatogenesis and is under positive selection. Table 6 shows that there are many protein coding genes present in these heterozygous regions. The RHZ on Chr6 is shared by every regional group but the Khoe and San. It has a length of 4 Mb, and has more than 140 protein coding genes including many members of the HLA complex family, olfactory receptor family, MHC class I genes, lymphocyte antigen 6 family, and the psoriasis susceptibility 1 candidate gene among others. As for ROHi, multiple RHZ are shared by different regional groups.

**Table 6.** *Location, length, percentage of individuals with ROH for the ROH island and protein coding genes of the three longest RHZ according to populations from global geographic regions*

|  | Chr | Pos 1 | Pos 2 | Length (Mb) | % Ind in ROH | Protein coding genes |
| --- | --- | --- | --- | --- | --- | --- |
| **Africa W.** |  |  |  |  |  |  |
|  | 6 | 287E+05 | 326E+05 | 4 | 1.5 | + 140 genes |
|  | 5 | 686E+05 | 711E+05 | 2.6 | 0.24 | <u>SLC30A5</u>, <u>ANP32A</u>, <u>CORO2B</u>, <u>GLCE</u>, <u>KIF23</u>, <u>LARP6</u>, <u>NOX5</u>, <u>PAQR5</u>, <u>RPLP1</u>, <u>TLE3</u>, <u>UAUCA</u>, <u>SPESP1</u>, <u>THAP10</u>, <u>THSD4</u> |
|  | 16 | 844E+05 | 869E+05 | 2.6 | 2.3 | <u>FOXL1</u>, <u>FOXC2</u>, <u>COTL1</u>, <u>COX4I1</u>, <u>CRISPLD2</u>, <u>EMC8</u>, <u>FAM92B</u>, <u>FOX</u>, <u>GINS2</u>, <u>GSE1</u>, <u>IRF8</u>, <u>KIAA0513</u>, <u>KLHL36</u>, <u>MTHFSD</u>, <u>TDLC1</u>, <u>USP10</u>, <u>AZDHHC7</u> |
| **Africa GG.** |  |  |  |  |  |  |
|  | 6 | 287E+05 | 326E+05 | 4 | 0.35 | + 140 genes |
|  | 12 | 1286E+05 | 1312E+05 | 2.7 | 2.8 | <u>ADGRD1</u>, <u>FZD10</u>, <u>GLT1D1</u>, <u>PIWIL1</u>, <u>RAN</u>, <u>RIMBP2</u>, <u>STX2</u>, <u>TMEM</u>, <u>SLC15A5</u> |
|  | 5 | 686E+05 | 711E+05 | 2.6 | 0 | See Africa W. Second RHZ |
| **Africa E.** |  |  |  |  |  |  |
|  | 6 | 287E+05 | 326E+05 | 4 | 0.51 | + 140 genes |
|  | 9 | 393E+05 | 428E+05 | 3.6 | 0 | <u>SPATA31A1</u>, <u>FOXD4L6</u>, <u>CBWD6</u>, <u>ANKRD20A2</u>, <u>CNTNAP3B</u> |
|  | 12 | 1278E+05 | 1312E+05 | 3.5 | 2.1 | See Africa GG. Second RHZ |
| **Africa H.** |  |  |  |  |  |  |
|  | 6 | 287E+05 | 326E+05 | 4 | 0.42 | + 140 genes |
|  | 12 | 1286E+05 | 1312E+05 | 2.7 | 2.8 | See Africa GG. Second RHZ |
|  | 5 | 686E+05 | 711E+05 | 2.6 | 0 | See Africa W. Second RHZ |
| **Africa S.** |  |  |  |  |  |  |
|  | 6 | 287E+05 | 326E+05 | 4 | 0.54 | + 140 genes |
|  | 9 | 388E+05 | 428E+05 | 4.1 | 0.2 | See Africa E. Second RHZ |
|  | 16 | 844E+05 | 869E+05 | 2.6 | 2.3 | See Africa W. Third RHZ |
| **Africa KS.** |  |  |  |  |  |  |
|  | 9 | 392E+05 | 428E+05 | 3.7 | 0.2 | See Africa E. Second RHZ |
|  | 8 | 64E+05 | 81E+05 | 1.8 | 3.3 | + 30 genes |
|  | 5 | 69E+06 | 706E+05 | 1.7 | 0 | See Africa W. Second RHZ |
| **Mix A.A.** |  |  |  |  |  |  |
|  | 6 | 287E+05 | 326E+05 | 4 | 0.4 | + 140 genes |
|  | 12 | 1278E+05 | 1312E+05 | 3.5 | 1.3 | See Africa GG. Second RHZ |
|  | 16 | 843E+05 | 873E+05 | 3.1 | 1.4 | See Africa W. Third RHZ |
| **Europe** |  |  |  |  |  |  |
|  | 9 | 388E+05 | 428E+05 | 4.1 | 0.03 | See Africa E. Second RHZ |
|  | 6 | 287E+05 | 326E+05 | 4 | 0.68 | + 140 genes |
|  | 15 | 202+E05 | 227+E05 | 2.6 | 0.35 | <u>GOLGA</u>, <u>OR4M2</u>, <u>OR4N4</u>, <u>POTEB2</u>, <u>POTEB3</u>, <u>LINC02203</u> |
| **Asia E** |  |  |  |  |  |  |
|  | 9 | 388E+05 | 428E+05 | 4.1 | 0.03 | See Africa E. Second RHZ |
|  | 6 | 287E+05 | 325E+05 | 3.9 | 0.42 | + 140 genes |
|  | 18 | 155+E05 | 185+E05 | 3.1 | 3.9 |  |
| **Mix H.A.** |  |  |  |  |  |  |
|  | 9 | 388E+05 | 428E+05 | 4.1 | 0.02 | See Africa E. Second RHZ |
|  | 6 | 287E+05 | 326E+05 | 4.0 | 0.3 | + 140 genes |
|  | 15 | 202+E05 | 227+E05 | 2.6 | 1.2 | See Europe. Third RHZ |

It is possible to use differences in ROHi and RHZ across regional groups to obtain a genetic distance that could provide an evolutionary perspective of the distribution of these homozygous and heterozygous genomic regions. Figure 9 shows a pairwise comparison of unique ROHi (A) and RHZ (B) in two heatmaps and, on the right of the figure, a rooted dendrogram for each heatmap using the percentage of unique RHOi or RHZ as genetic distances. Both rooted dendrograms present similarities and differences in their branching. Both establish two main groups: SSA and out-of-Africa. Within SSA (with the exception of the Horn of Africa), both dendrograms first split off the Khoe and San from the rest of groups and then both split Bantu-speaking populations from Southern Africa from the rest. Also, both dendrograms, include the mixed African-American group in the SSA branch. In the out-of-Africa branch both dendrograms group together European and admixed Hispanic-American populations. The biggest differences between the two dendrograms is where they locate the Horn of Africa populations; the ROHi dendrogram groups them with the out-of-Africa branch, whereas the RHZ dendrogram groups them with the SSA branch.

**Figure 9.**
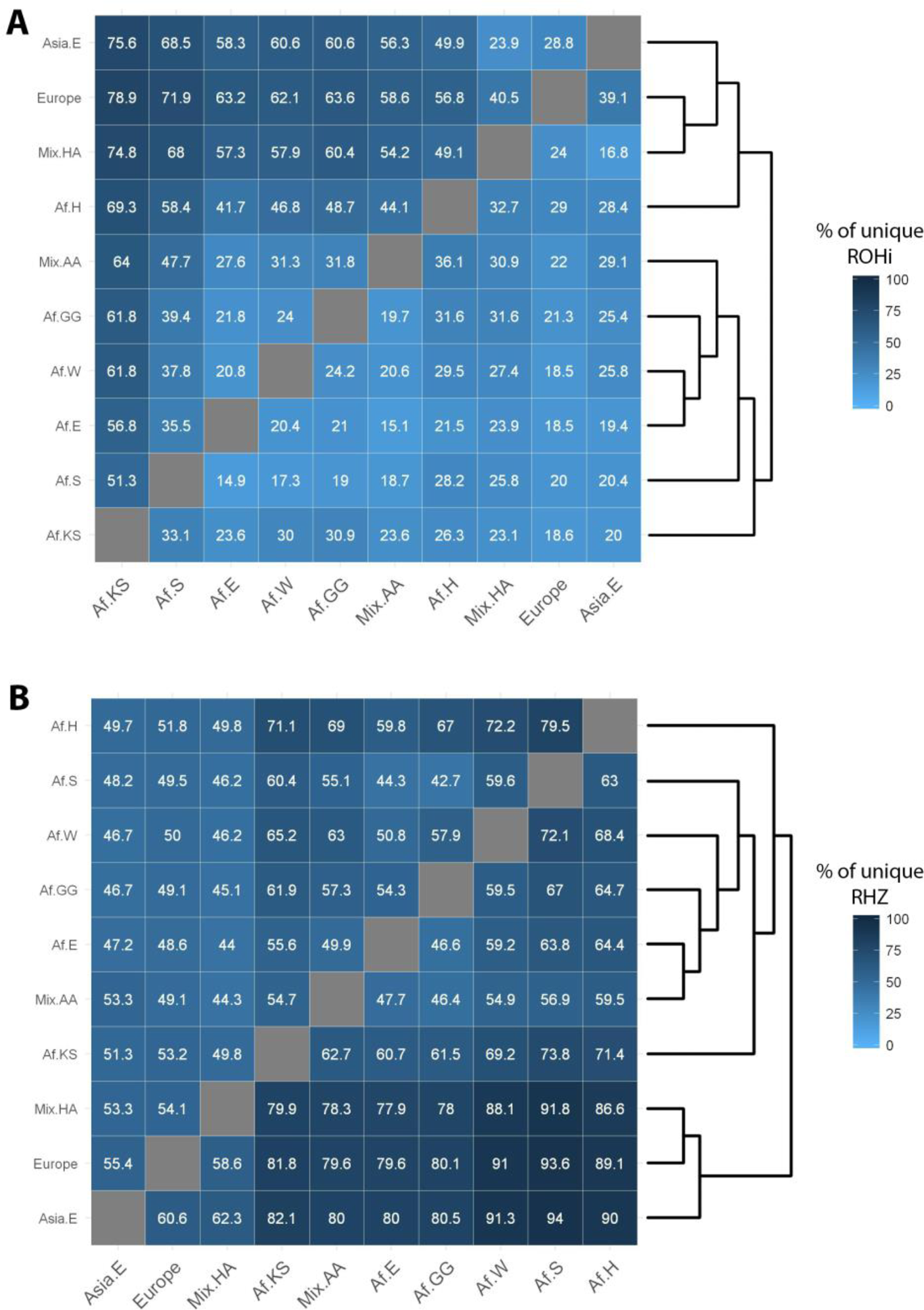
Heatmap and rooted dendrogram of the unique ROH islands (A) or RHZ (B) per geographical regional group and admixed populations. The heatmap shows pairwise % of unique ROHi/RHZ between regional groups. The rooted dendrogram was obtained using optimal leaf ordering or OLO. Af.KS: African Khoe and San populations; Af.S: population from southern Africa; Af.E: population from eastern Africa; Af.W: population from western Africa; AF.GG: population from the Gulf of Guinea; Mix.AA: African-American admix populations; Mix.HA: Hispanic-American admix populations; Europe: European populations; Asia.E: populations from eastern Asia.

## DISCUSSION

SSA populations have been the subject of extensive genomic research with the objective of understanding their demographic history, current population structure, selection footprints and to advance the field of biomedical genetics^2-4; 62-65^. To achieve these objectives classic population structure tools like F_ST_, admixture analysis, and PCA are often used. ROH analyses have not yet been fully explored even though their usefulness as a tool to decipher different demographic histories is clear and studies range from research on individuals to describing elaborate worldwide population-based trends^7; 16^. For example, we have shown (Figures 2 and 3) that populations around the globe experience a reduction in the mean total length of ROH in length categories above 0.5Mb. Since the length of ROH is inversely proportionate to its age, a possible explanation for this global phenomenon could be that populations around the world experienced a size increase about the same time, reducing autozygosity provoked by low N_e_ and genetic drift. However, to put these results into context and compare them to the estimates of population size already published^27; 66^, it is necessary to determine the age of the different ROH sizes. Preliminary results estimate that ROH length of 1.5Mb may have a median age of approximately 30 generations (*personal communication D.W. Clark*) and ROH longer than 4 Mb may not be older than 10 generations^8^.

Previous studies in SSA showed that Africa is the continent with the smallest burden of ROH and that within Africa there is limited heterogeneity in ROH distribution, occurring essentially between the hunter-gatherers and the agro-pastoralists^7; 20; 23^. Our study, however, shows that ROH distribution in SSA is very heterogenous and much more complex than expected, with different scenarios for ROH shorter and longer than 1.5Mb. Although the vast majority of SSA populations have a low burden of short ROH, that is not the case for long ROH where we find SSA populations with a higher burden in comparison to other populations around the globe. In contrast with previous studies, our fine scale analysis has overcome some limitations: It has representation of populations from Western, Eastern and Southern Africa; it uses high-density SNP coverage (~1.2 M SNPs after QC) providing good resolution to accurately call for ROH; the PLINK software conditions for ROH calling were optimized to accurately call short ROH; and analyses were developed to understand the ROH distribution and its demographic consequences.

### Insights into the past - analysis of short ROH (ROH<1.5Mb)

The demographic history of SSA is characterized by large effective population sizes over many generations that have led to high genetic diversity, shorter LD structures and lower burden of small ROH^;24^. Our study reports considerable structure in the distribution of short ROH in Africa with populations from the Horn of Africa (Somali, Oromo and Amhara) having the largest burden of ROH <1.5Mb. In the absence of evidence to support a different evolutionary trajectory of the effective population size between these and other SSA populations, the most plausible explanation is that the short ROH were introduced through admixture of Semitic and Cushitic populations with others from the Arabian Peninsula. It has been found that Ethiopian individuals are characterized by a large (40-50%) non-African genetic component most likely originating mainly from Egypt, the Levant and Yemen in a migration that took place approximately 3 thousand years ago (Kya)^28; 67^. This hypothesis is also supported by the ROHi profiling of populations in the Horn of Africa that have the highest number of short ROHi (0.1 – 0.3Mb) and the shortest mean ROHi length (0.37Mb) (Table 3), with 83% of ROHi shorter than 0.5Mb. When compared with other regional groups (Figure 9), the populations from the Horn of Africa share more ROHi with regional groups outside Africa (Figure 9A). There is a reasonably homogeneous burden of short ROH between Western, Gulf of Guinea, Eastern and Southern Bantu-speaking groups (Table 1 and Figure 4), but the Khoe and San, having split from non-Khoe and San lineages 100 to 150 Kya^68^, show heterogeneity (e.g. Northen Ju, Ju/’hoansi have a similar burden to populations in Western Africa, and the Central Khoe-Kwadi and Khwe, have the lowest burden in all SSA).

The shape of the distribution of the ROH <1.5Mb shown in Figure 4 is also highly informative. Admixed populations, originating from ancestral populations with different ROH burden, would have individuals with different Sum of ROH<1.5Mb due to their distinct coalescent histories, as is shown in Figure 4 where most of the admixed populations present platykurtic and skewed distributions. Hispanic-American populations (CLM, PUR, MXL), with ROH<1.5Mb burden similar to Europeans have a small proportion of African ancestry (7.8%, 13.9% and 4.3% respectively) but higher proportion of European (66.6%, 73.2% and 48.7% respectively) and Native American (25.7%, 17.9% and 47.0% respectively) ancestry^69; 70.^ The PEL population has shorter ROH due to a greater Native American ancestry (2.5% African, 20.2 European and 77.3% Native American)^69; 70^. For these populations ROH<1.5Mb arose before the time of admixture; estimated as 14 generations for CLM, 7 for MXL and 16 for PUR. PEL population was found to have two different admixture pulses 12 and 5 generations ago, with the last one being 91.1% Native American^70^. On the opposite side, African-American admixed populations (ASW and ACB) have reasonably normal distributions with almost no skewness. These two populations seem to have a very tight distribution and small burden of ROH<1.5Mb, similar to the Western Africans and Guinea Gulf populations. This could be explained by the elevated proportion of African ancestry (88% and 75.6% respectively) and small proportions of European and Native American ancestry (ACB: 11.7% European, 0.3 Nat American; ASW: 21.3% European, 3.1% Nat American)^69; 70^. The South African Coloured populations, another example of recently (150-300 years) highly admixed populations, have a ROH<1.5Mb burden very similar to Khoe and San populations. Nevertheless, different studies reported different ancestry components for Coloured populations arising from Khoe, San, and Bantu speakers, as well as European, South Asian and Austronesian populations ^6; 71^ giving insight into the complexity of these admixed populations. Finally, it is also possible to detect kurtosis and skewness in some Khoe and San populations which would indicate admixture. Unequivocally, /Gui//Gana, Nama, Karretjie and ≠Khomani distributions for sum of ROH<1.5Mb reveal their admixture origins. In these four Khoe and San populations Bantu and even European ancestral components were found^23; 72; 73^.

### Consanguineous cultural practices and modern genetic isolation - analysis of long ROH (ROH>1.5Mb)

The study of ROH>1.5Mb is very useful to shed light on the role of cultural practices in genome homozygosity levels. Different anthropological and human biology studies have systematically identified African populations with a clear cultural preference for consanguineous marriages, and some that purposely avoid such unions^74-83^. For example, one of the most recently published studies, which analysed 548 marriages over the period 1994-96 in the Fulani from Burkina Faso, found that 399 marriages (68.3%) were between relatives and 185 (31.7%) were between non-related individuals. The average inbreeding coefficient (α) was estimated as 0.0364^82^. Similar inbreeding coefficients were found by other studies, for example an α=0.0322 in the Khartoum population from Sudan^79^. Our study shows a very heterogeneous distribution of ROH>1.5Mb among SSA: populations with very little burden of long ROH>1.5Mb, and completely absence of ROH>4Mb, for example in the Amhara from the Horn of Africa, the Yoruba from the Gulf of Guinea or the Kikuyu from Eastern Niger-Congo Africa, and populations with a high burden of ROH>1.5Mb like the Somali from the Horn of Africa, the Fula from Western Africa or the Khoe and San !Xun and Ju/’hoansi. A heterogeneous distribution of long ROH was found within SSA regions: Somali and Oromo populations, from the Horn of Africa, speak Cushitic languages, but Somalis are predominantly Sunni Muslims, with a preference for first-cousin unions, while Oromo people are predominantly Ethiopian Orthodox or follow traditional religions with no preference for consanguineous unions^84^. Despite the results presented in this study, in other SSA regions like Guinea Gulf or Eastern Africa anthropological studies there are groups with cultural preferences for unions between relatives like the Futajalonke from Guinea^83^, the Baoule from Ivory Coast^83^, the Ewe from Ghana^83^, Arab groups in Kenya^77^, the Kigali and Tutsi from Rwanda^74^ and the Khartoum and Gezira groups from Sudan^79^. Cultural differences among individuals within populations can be inferred from the shapes of the distributions in Figure 4. Not surprisingly, populations with larger burden of ROH>1.5Mb (in order: !Xun, Ju/’hoansi, Somali, Khew, PEL, Gui//Gana, CLM, Fula, etc.) have the longest right tails and the highest number of individuals with an inbreeding coefficient higher than F=0.0152 (Figure 5). Hence, despite previous reports, we have found African populations with mean genomic inbreeding coefficients (F_ROH_) higher than several other isolated populations around the world, such as the PEL from Lima in Peru.

In order to sketch a more complete picture of genomic homozygosity in SSA populations, it is important to analyse the origins of this homozygosity. The representation of the mean number of ROH compared to the mean total sum of ROH showed a right shift for Khoe and San populations like Ju/’hoansi, !Xun, Gui//Gana or Khwe, indicating the possible presence of recent consanguineous loops and a deviation from panmixia (Figure 6A). However, if the influence of the F_IS_ in the F_ROH_ is represented as shown in Figure 6B, a different picture is revealed. In summary, it is possible to establish a classification with 3 main groups characterized by demographic history. Firstly, populations with different levels of cultural consanguinity practices like Somali, Fula, CLM, GIH and Wolof. Secondly populations with low levels of inbreeding provoked by their large continental N_e_, in this group we can find the bulk of Europe, Asian and SSA populations. Thirdly, populations with considerable genetic drift and recent genetic isolation like PEL, Khwe, Ju/’hoansi, !Xun and Herero. The representation of F_IS_ vs F_ROH_ is a better approach to identify the origins of inbreeding since it provides information about the proportion of F_ROH_ due to deviation from panmixia or from genetic isolation. Furthermore, this representation is helpful to identify populations with an excess of homozygotes possibly due to the Wahlund effect, which may be expected for the Gui//Gana population, or, more surprisingly, with the Southern Tuu-speaking Khoe and San, the ≠Khonami and Karretjie peoples.

### Genomic distribution of ROH and the identification of regions under selection

Examining ROH has been shown to be useful for studying genome biology and to identify regions under selection^19-21^. The existence of ROH islands (ROHi) and regions of heterozygosity (RHZ) can be explained in part as a consequence of stochastic processes across the genome, or by variation of the effects of demographic processes across the genome, influencing genetic diversity^7; 20^. However, there is increasing evidence that ROH islands may be a consequence of positive selection processes that reduce haplotype diversity and increase homozygosity around the target locus, increasing ROH frequencies in the regions under selection^20; 85^. Besides the presence of specific protein coding genes, previously detected to be under positive selection, in the five most prevalent ROHi (Table 4 and 5), we identified other genes previously shown the be under positive selection in African populations^2; 23; 65^. Different loci associated with infectious disease susceptibility and severity, including *HP*^2^, *CLTA4*^86^ and *PKLR*^87^ for malaria, *IFIH1*^88^ and *OAS2*^2^ for Lassa fever, *FAS*^89^ for Trypanosomiasis and other genes involved in general immune response (e.g. *PRSS16*^23^ and *POM121L2*^23^) were found within ROHi in different geographical regions. For example, *CTLA4* was found in ROHi in every region, but *HP* and *PKLR* were found to be in ROHi just in Western and Eastern SSA and in the Horn of Africa. Other genes related to trypanosomiasis infection and kidney disease, like *APOL1*^90^, or to different forms of hypertension, like *ATP1A1*^2^, *AQP2*^2^ and *CSK*^2,91^ were found in ROHi in different regions from SSA. As was shown in Table 6 within RHZ haplotypes it is also possible to find multiple protein coding genes related to diverse biological functions like immune response (*HLA* complex or *IRF* gene family), cellular cycle (*ANP32A*^92; 93^), chromosomal aberrations (like different members of the *GOLGA* gene family^94^) cancer (*NOX5*^95^), brain development (*KIAA0513*^96^) and olfactory receptors (*OR* gene family) among others. These heterozygous regions might represent haplotypes enriched for variants that have a negative impact on fitness in homozygosity, or regions that harbor loci with heterozygote advantage (overdominance) under any form of balancing selection. Furthermore, this hypothesis is also supported by the fact that it is possible to establish differences and similarities between the locations of ROHi and RHZ between populations from different geographic regions, as it is shown in Figure 9. Furthermore, since the majority (more than 75%) of ROHi and RHZ identified in this study include genomic regions that had previously been identified as sites of recent selection, this analysis raises the possibility that other loci in ROHi and RHZ may also harbor genes that have been subjected to positive or balancing selection.

## Conclusion

Detailed ROH analysis demonstrated a heterogeneous distribution of autozygosity across SSA populations shedding light on the complex demographic history of the region. While short ROH (ROH<1.5Mb) provided insights into effective population size and past admixture events, long ROH (ROH>1.5Mb) informed us about the impact of consanguineous cultural practices, modern endogamy and genetic isolation. We also showed that ROHi and RHZ can be used to identify genomic regions under selection pressure. Studying a better representation and larger sample size across different SSA populations will provide more nuanced interpretations of demographic histories. The H3Africa (Human Heredity and Health in Africa) initiative is generating genomic data including whole genome and exome sequences and genome-wide genotyping using an African tailored array that captures common genetic diversity in African genomes^3; 4^. The added value of this resource lies in its rich phenotype and clinically relevant data that will enable biomedical research across the continent making it possible to study the distribution of ROH and RHZ in common complex traits.

## Supplemental Data

Supplemental Data include eight figures and Supplemental Material and Methods including the optimization of PLINK ROH calling algorithm to obtain short ROH and the comparison of ROH obtained from the same samples with different SNP coverage.

## Acknowledgments

FCC is a National Research Foundation of South Africa (NRF) postdoctoral fellow and MR holds a South African Research Chair in Genomics and Bioinformatics of African populations hosted by the University of the Witwatersrand, funded by the Department of Science and Technology and administered by the NRF.

## Footnotes

^1^ The term *Khoe-San* is often used in the literature, but is regarded by some as offensive as it conflates two distinct groups. The impact of colonialism had a very traumatic effect on population size and structure. We use the phrase *Khoe and San* to describe people who have either Khoe and/or San ancestry as a neutral term to describe people who live in similar regions and have had some shared history in the last centuries.

## Supplemental Material and Methods

### Description of the Data and the Methodology

PLINK’s observational approach underestimates small ROH (shorter than 500Kb) when using recommended conditions (50 as the minimum number of SNP that the PLINK’s sliding window, and ROH, is required to have) in array-genotyped data in comparison to whole genome sequence low coverage^1^. For the analysis of the current study it is important to have accurate ROH estimates for sizes as short as 300 Kb. In order to achieve this goal, we tested different PLINK parameters of ROH calling in array-based data and compared them with ROH obtained from low coverage (3-6x) whole genome sequence. We therefore published the required PLINK conditions to obtain equivalent results, with parameters for ROH longer than 1.5Mb, between WGS low coverage and SNP array technologies^1^. In the current study we used the same conditions as a starting point to obtain equivalent short ROH estimations.

Individuals with both genome-wide SNP genotypic data and WGS low coverage data from the 1000 Genomes Project – Phase 3 (KGP) and the African Genome Variation Project (AGVP) were used. For both datasets the Infinium Omni 2.5-8 Bead chip from Illumina was used. The KGP includes a total of 1685 individuals from 18 populations with genotypic data available from array and WGS low coverage (4x): European ancestry FIN (n=99), GBR (n=91), IBS (n=105), TSI (Tuscani n=102) and CEU (n=99); African-American ancestry ASW (n=61) and ACB (n=96); Hispanic-American ancestry PUR (n=104), PEL (n=85), CLM (n=95) and MXL (n=100); Eastern Asia ancestry CDX (n=98), CHB (n=100), CHS (n=105), JPT (n=100) and KHV (n=99); and African ancestry YRI (n=108) and LWK (n=99). The AVGP includes 200 samples (100 Zulu and 100 Baganda) where array-genotype data and WGS low coverage (4x) are available. For each population, data from both array genotyping and WGS were filtered to remove MAF <0.05 and those diverging from H-W with p <0.001. Only SNPs of the 22 autosomes were included in the analysis.

We used PLINK v1.9 to identify ROH. The following conditions were used to call ROH in the WGS low coverage data`--homozyg-snp 50, --homozyg-kb 300, --homozyg-density 50, --homozyg-gap 1000, --homozyg-window-snp 50, --homozyg-window-het 3`. For array-genotype data the following conditions where used: `--homozyg-snp (30, 40, 50), --homozyg-kb 300, --homozyg-density (30, 40, 50), --homozyg-gap 1000, --homozyg-window-snp (30, 40, 50), --homozyg-het 1.`

Using violin plots for visualisation of the ROH data distribution, we performed an exploratory data analysis comparing five different ROH class sizes obtained from array-genotype and WGS data. Class 1: 300Kb<ROH≤500Kb; Class 2: 500Kb<ROH≤700Kb; Class 3: 700Kb<ROH≤900Kb; Class 4: 900Kb<ROH≤1000Kb; Class 5: 1000Kb<ROH≤1500Kb.

### Results and Conclusions

In Figures S1 to S5 show violin plots of the sum of ROH for the five classes of ROH lengths. For each of the continental divisions (Africa: Figure S1; Hispanic-American: Figure S2; African-American: Figure S3; Asian: Figure S4 and Europe: Figure S5) we demonstrate that some adjustments are appropriate when dealing with array-genotype data. For example, when we relax PLINK’s conditions to 30 SNPs per sliding window and ROH, it is possible to obtain more equivalent sum of ROH estimates for Class 1 and 2 (300Kb to 700Kb) than when using previously recommended conditions (50 SNP). Furthermore, the sum of ROH estimates didn’t change much when considered ROH longer than 700Kb.

According to these results we can conclude that by using a sliding window of 30 SNPs in PLINK we can obtain a better estimation of short ROH that does not interfere with the estimation of longer ROH.

## Supplemental References

1. Ceballos, F.C., Hazelhurst, S., and Ramsay, M. (2018). Assessing runs of Homozygosity: a comparison of SNP Array and whole genome sequence low coverage data. BMC Genomics 19, 106.

**Figure S1.**
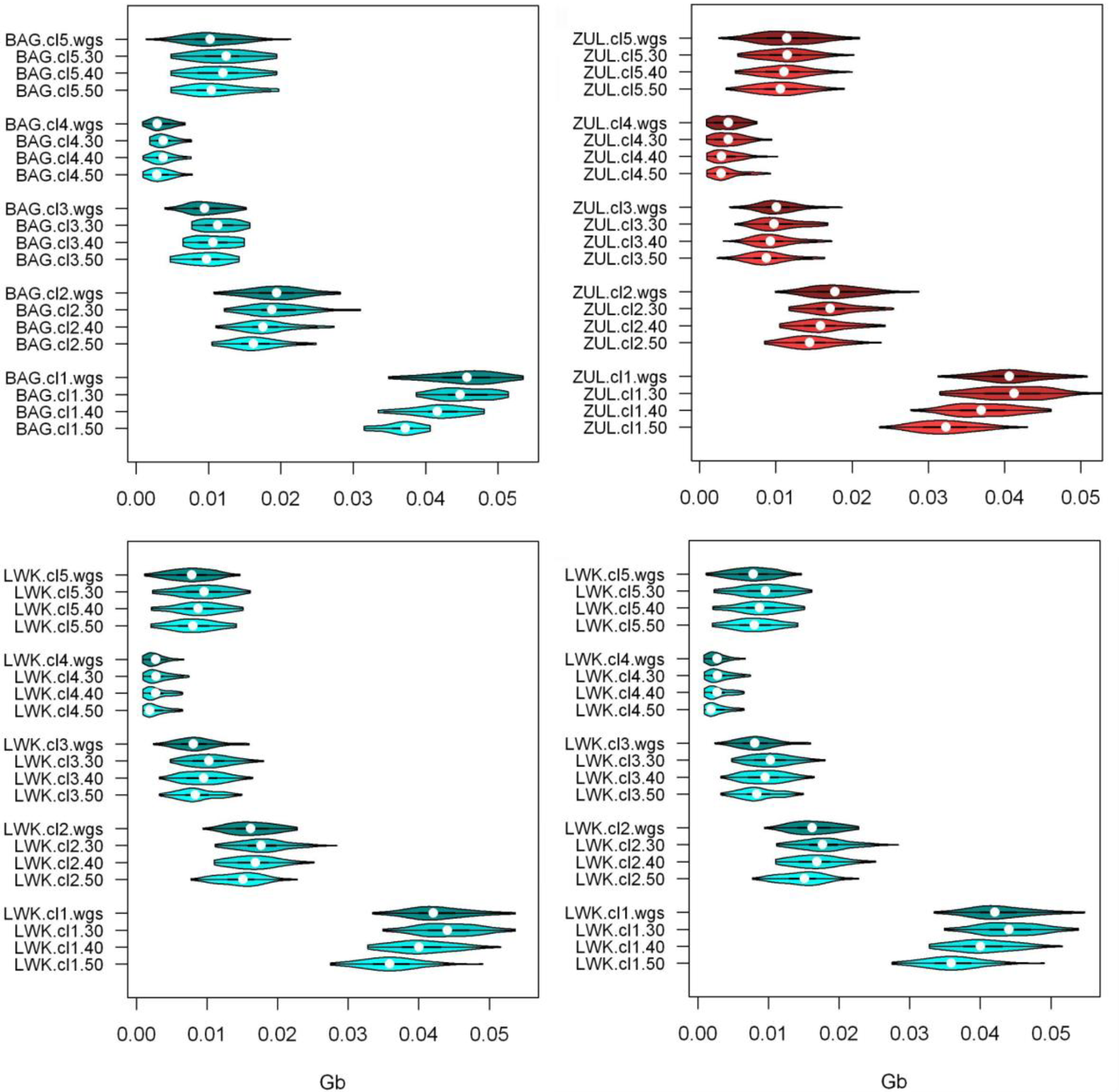
Violin plots of the sum of ROH for 5 classes of ROH length in African populations from 1KGP and AGVP with Array and WGS data available Cl1: 0.3Mb<ROH≤0.5Mb; Cl2: 0.5Mb<ROH≤0.7Mb; Cl3: 0.7Mb<ROH≤0.9Mb; Cl4: 0.9Mb<ROH≤1.0Mb; Cl5: 1Mb<ROH≤1.5Mb. BAG: Baganda population from AGVP; ZUL: Zulu population from the AGVP; LWK: Luhya population from the 1KG; YRI: Yoruba population from the 1KGP. For each population, 30, 40 and 50 SNPs per window as PLINK conditions to obtain ROH with the Array data were compared with ROH from WGS data by using a window of 50 SNPs

**Figure S2.**
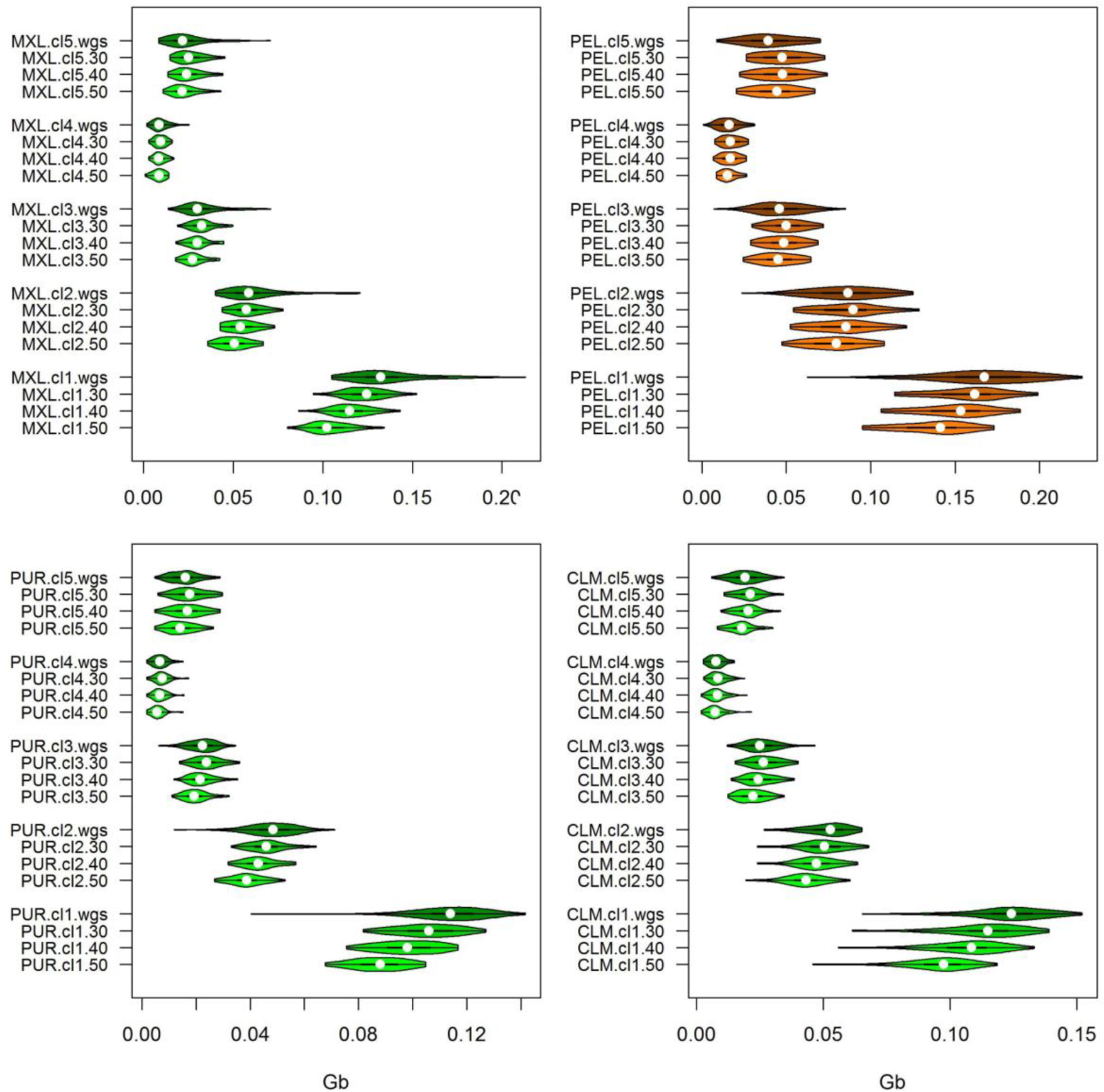
Violin plots of the sum of ROH for 5 classes of ROH length in American populations from 1KGP with Array and WGS data available. Cl1: 0.3Mb<ROH≤0.5Mb; Cl2: 0.5Mb<ROH≤0.7Mb; Cl3: 0.7Mb<ROH≤0.9Mb; Cl4: 0.9Mb<ROH≤1.0Mb; Cl5: 1Mb<ROH≤1.5Mb. For each population, 30, 40 and 50 SNPs per window as PLINK conditions to obtain ROH with the Array data were compared with ROH from WGS data by using a window of 50 SNPs.

**Figure S3.**
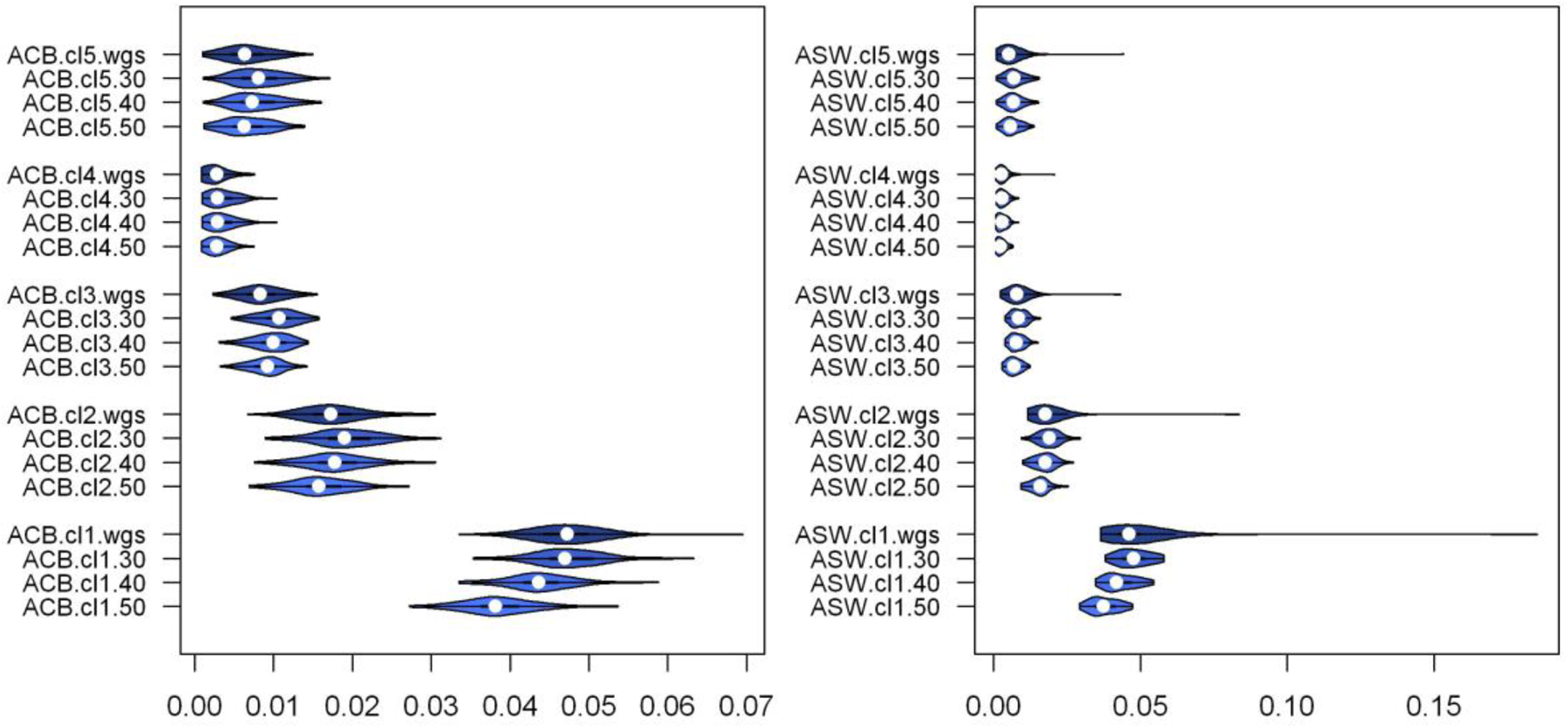
Violin plots of the sum of ROH for 5 classes of ROH length in admixed African - American populations from 1KG with Array and WGS data available. Cl1: 0.3Mb<ROH≤0.5Mb; Cl2: 0.5Mb<ROH≤0.7Mb; Cl3: 0.7Mb<ROH≤0.9Mb; Cl4: 0.9Mb<ROH≤1.0Mb; Cl5: 1Mb<ROH≤1.5Mb. For each population, 30, 40 and 50 SNPs per window as PLINK conditions to obtain ROH with the Array data were compared with ROH from WGS data by using a window of 50 SNPs.

**Figure S4.**
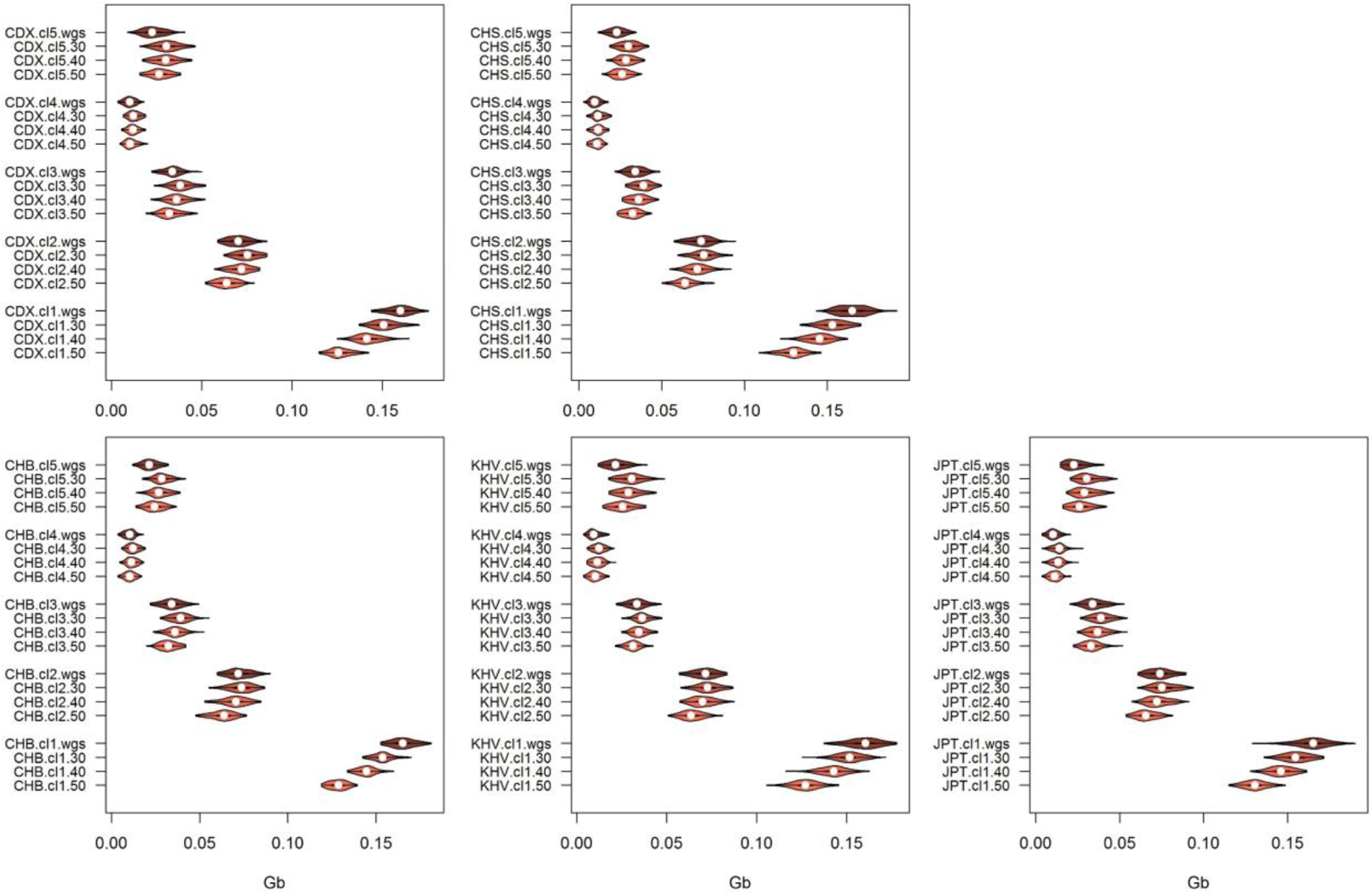
Violin plots of the sum of ROH for 5 classes of ROH length in Eastern Asia populations from 1KGP with Array and WGS data available. Cl1: 0.3Mb<ROH≤0.5Mb; Cl2: 0.5Mb<ROH≤0.7Mb; Cl3: 0.7Mb<ROH≤0.9Mb; Cl4: 0.9Mb<ROH≤1.0Mb; Cl5: 1Mb<ROH≤1.5Mb. For each population, 30, 40 and 50 SNPs per window as PLINK conditions to obtain ROH with the Array data were compared with ROH from WGS data by using a window of 50 SNPs.

**Figure S5.**
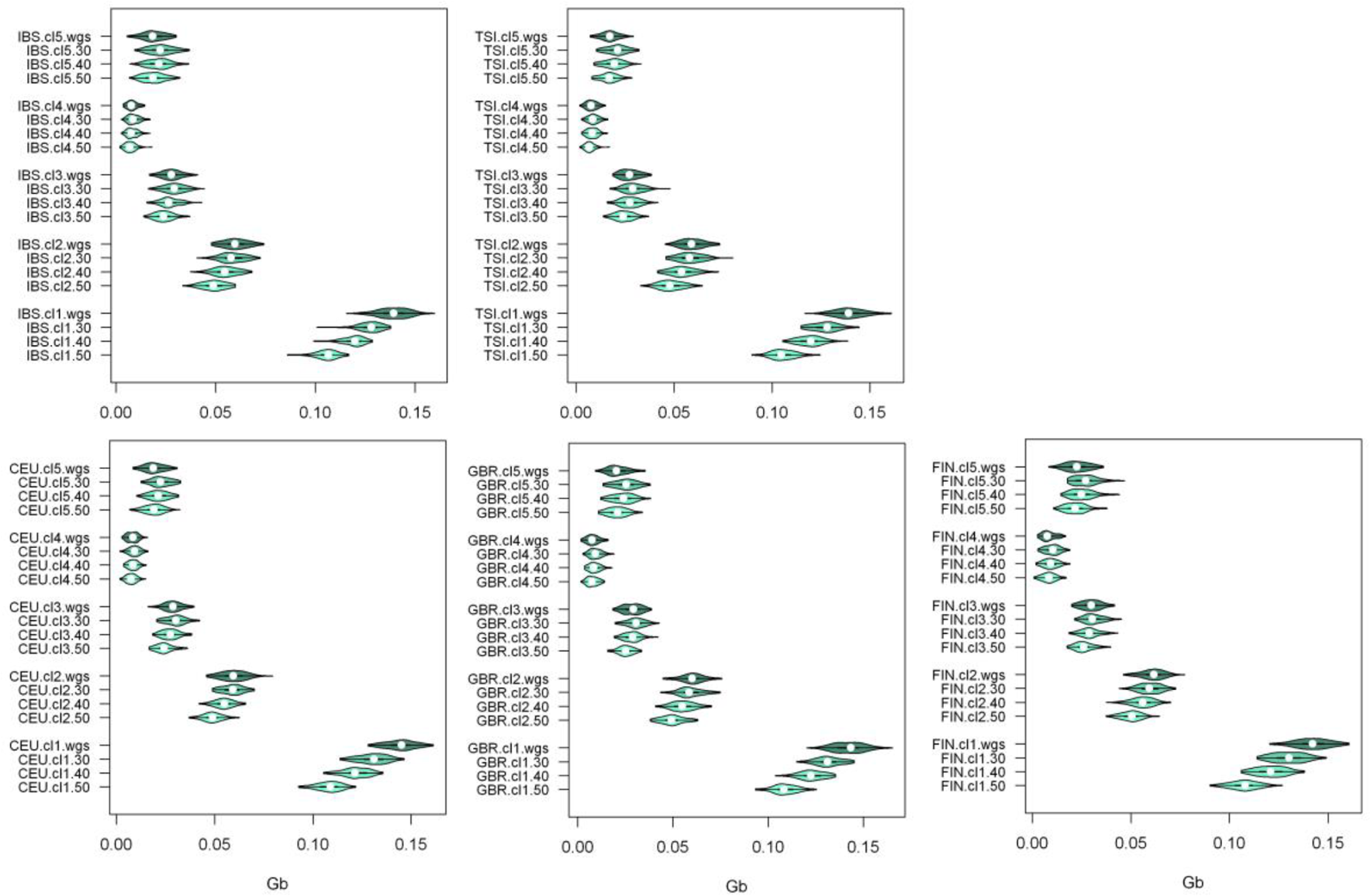
Violin plots of the sum of ROH for 5 classes of ROH length in European populations from 1KG with Array and WGS data available Cl1: 0.3Mb<ROH≤0.5Mb; Cl2: 0.5Mb<ROH≤0.7Mb; Cl3: 0.7Mb<ROH≤0.9Mb; Cl4: 0.9Mb<ROH≤1.0Mb; Cl5: 1Mb<ROH≤1.5Mb. For each population, 30, 40 and 50 SNPs per window as PLINK conditions to obtain ROH with the Array data were compared with ROH from WGS data by using a window of 50 SNPs

**Figure S6.**
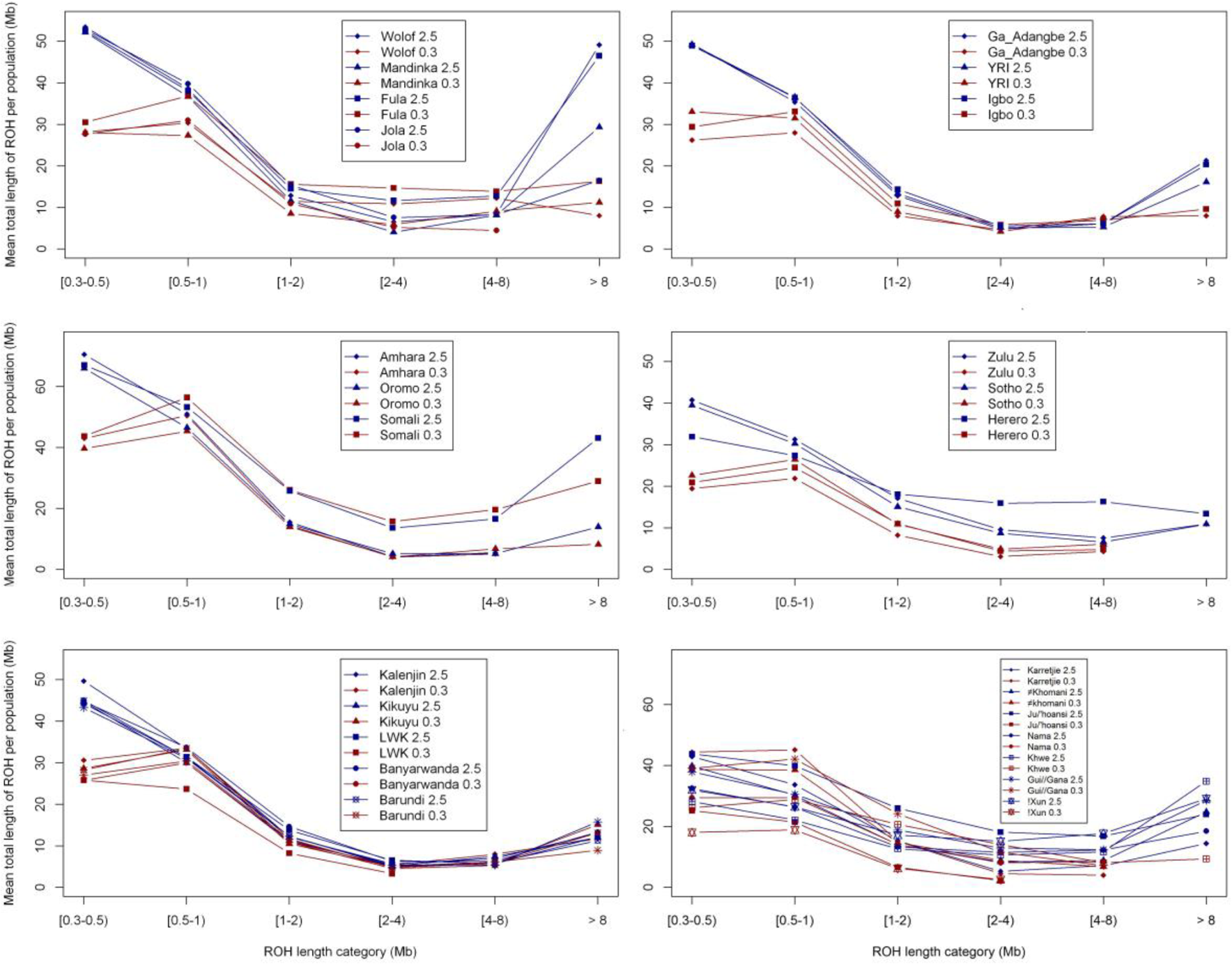
Mean total sum of ROH in different length categories. Blue colored lines represent the populations not being merged (Array of 2.5 M SNPs). Red colored lines represent the outcome of the different datasets (AGVP, Schlebusch et al. 2012, KGP, HGDP) after being merged (382,840 SNPs available).

**Figure S7.**
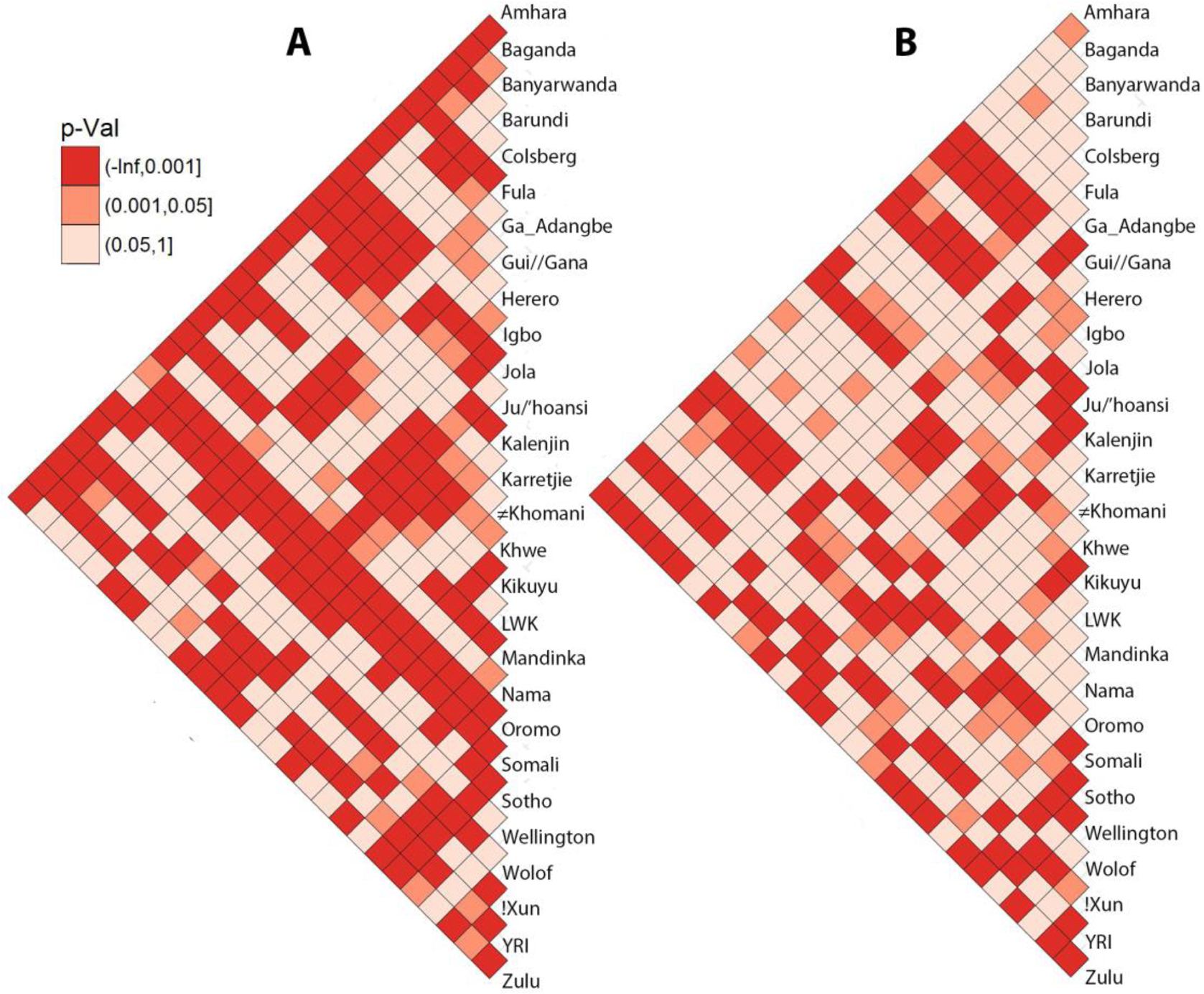
Pairwise comparisons of populations within Sub-Saharan Africa by the Mann-Whitney-Wilcoxon non-parametrical test (MWW) of ROH shorter than 1.5Mb (A) and ROH longer than 1.5Mb (B).

**Figure S8.**
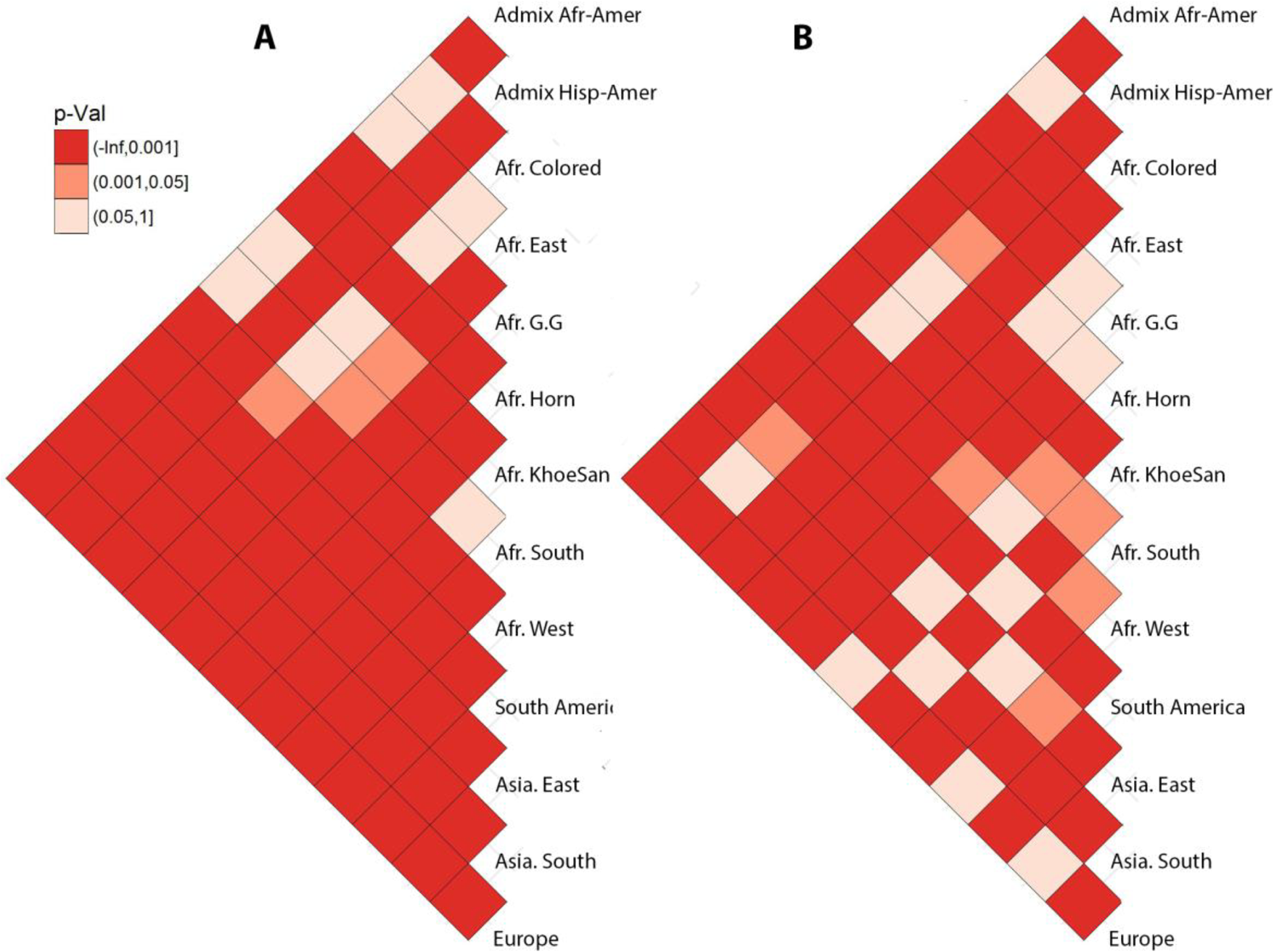
Pairwise comparisons of regional groups by the Mann-Whitney-Wilcoxon non-parametrical test (MWW) of ROH shorter than 1.5Mb (A) and ROH longer than 1.5Mb (B).

